# Know Thy Neighbour: Benchmarking Protein Language Model Embedding-Based Annotation Reveals Anisotropy Substantially Impacts Retrieval

**DOI:** 10.64898/2026.11.13.766344

**Authors:** George Bouras, Robert A. Edwards

**Affiliations:** School of Medicine, College of Health, Adelaide University, Adelaide, SA 5005; College of Science and Engineering, Flinders University, Bedford Park, SA 5042

## Abstract

Functional annotation using protein language models (‘pLMs’) usually proceeds by transferring the function of a query’s nearest neighbour in a reference database. A major drawback is that embedding-based annotation transfer has no null model, i.e. no accepted way to set a cosine-similarity threshold to control false positives. This contrasts with traditional bioinformatics methods, including sequence- and structure-based alignment tools, which rank hits with E-values derived from explicit null models. Here, we benchmark final-layer, mean-pooled embeddings from twelve models against MMseqs2 and Foldseek for remote-homology retrieval on curated datasets based on Swiss-Prot and SCOPe, fixing per-model thresholds at a 1% false-positive rate and probing the score with a ladder of synthetic decoys. On Swiss-Prot, pLMs without retrieval training recover at most 15% of remote homologs at this operating point, against 85% for MMseqs2, and for the ESM-2 and ESM-C families shuffled decoys have *higher* cosine similarities to their nearest database hits than real remote homologs do. These two pLM families exhibit extreme anisotropy, using fewer than 15 effective dimensions out of 640–2560, and low effective dimensionality is strongly associated with poor performance across models. Applying ZCA whitening, fit on the reference database, to the embeddings sharply raises threshold-free Top-1 superfamily retrieval, e.g. for ESM-2 650M from 59% to 88% on SCOPe, and generally improves remote homology sensitivity at a fixed false-positive rate. As correcting embedding anisotropy requires only the reference database and is compute cheap, it should be considered and tested for any pLM embedding-based annotation task.

## 1 Problem

### 1.1 Background and Traditional Bioinformatics Approaches

Protein language models (‘pLM’) are (mostly) transformer-based language models trained on a large corpus of protein sequences [Elnaggar et al., 2022, Lin et al., 2023]. Although usually trained with masked or autoregressive objectives using only amino acid sequence, pLM embedding space has been shown to capture elements of protein function [Jha et al., 2025] and structure [Zhang et al., 2024].

Protein functional annotation has traditionally been conducted using sequence homology-based approaches. Such methods include alignment-based tools such as BLAST [Altschul et al., 1990] and its successors DIAMOND [Buchfink et al., 2015] and MMseqs2 [Steinegger and Söding, 2017], which align query proteins against a reference database of proteins with annotated functions.^1^ More recently, the advent of AlphaFold2 [Jumper et al., 2021], along with fast and sensitive protein structural aligners such as Foldseek [van Kempen et al., 2024] (which tokenises a protein’s 3D coordinates into a 1D representation known as Foldseek ‘3Di’) has allowed for structure homology-based annotation utilising similar algorithms to sequence homology-based methods. Usually, sequence- and structure-based annotation proceeds by transferring the reference database top hit function to the query if it passes certain significance thresholds. A major reason why sequence- and structure-based methods have proliferated is that these methods are calibrated using a null model, meaning that top scoring hits passing a reasonable E-value threshold can generally be trusted to be true positives. As an example, BLAST uses Karlin–Altschul statistics that give principled E-values conditioned on query length, amino acid composition and database size [Karlin and Altschul, 1990, Altschul et al., 1997]. Structure-based methods take a similar approach [van Kempen et al., 2024]. While there have been critiques of the calibration and limits of both sequence- [Lu et al., 2024] and structure-based [Edgar and Sahakyan, 2025] E-values, their results are generally trustworthy and indeed are widely used for annotation across biological domains.

40th Conference on Neural Information Processing Systems (NeurIPS 2026). Workshop: I Can’t Believe It’s Not Better (ICBINB): Failure Modes of AI in Biology.

### 1.2 No Null Models For pLM Embeddings

The capacity of pLM embeddings to encode functional and structural information suggests that proteins in a similar area of the embedding space are likely to share similar structures and functions. Functional annotation transfer between proteins with homologous embeddings is known as embedding-based annotation transfer (‘EAT’) [Heinzinger et al., 2022]. The usual approach is to embed both the query and reference database proteins, mean-pool per-residue embeddings, retrieve the nearest neighbour below a certain Euclidean distance or above some cosine similarity threshold and then transfer the neighbour’s annotation [Littmann et al., 2021, Senoner et al., 2025, Jha et al., 2025]. This threshold is database and especially model specific. While EAT can find relationships between proteins in and beyond the twilight zone of sequence alignment below 35% sequence identity [Rost, 1999], the primary issue^2^ with its deployment into protein annotation workflows is the difficulty in calibrating thresholds that can discriminate true positives from true negatives. Existing methods and benchmarks generally report ranking-only evaluations in which every query has a true homolog, omitting queries that are true negatives [Jha et al., 2025, Heinzinger et al., 2022, Kabir et al., 2024, Banerjee et al., 2026]; on such metrics pLMs can even appear to outperform sequence alignment [Kabir et al., 2024]. Without a null model as used to calibrate E-values for sequence- and structure-based search, cosine similarity based scoring can be difficult to trust. There is therefore a benchmarking gap in assessing the calibration of EAT methods between different pLMs, and also in comparison to existing traditional bioinformatics methods.

## 2 Proposed approach

### 2.1 Datasets

Creating benchmark datasets for homology-based inference and protein function annotation is extremely difficult. Sequence- and structure-based methods themselves are generally used for dataset curation, including Test 1 (and implicitly, Test 2) below. Therefore, it is not necessarily expected that pLM-based methods outperform traditional methods, leading us to define answerable and remote as described below.

#### 2.1.1 Test 1 - Swiss-Prot/Pfam Sequence Based Splits

All 393,237 proteins with length up to 400 AA from UniProtKB/Swiss-Prot release 2026_02 [Consortium, 2023] were fetched via the UniProt REST API. These entries were filtered, keeping 370,558 proteins containing at least one Pfam family (based on v38.2) [Mistry et al., 2021] with only standard AA residues.

All sequences were then clustered using MMseqs2 v18.8cc5c [Steinegger and Söding, 2017] easy-cluster at 30% identity, 80% bidirectional coverage (--cov-mode 0). 20% of clusters were held out as queries, while the remaining 80% constitute the reference database. Overall, this yielded 73,751 initial queries and 296,807 reference proteins.

Each query protein was then denoted as *answerable* if the query shared at least 1 Pfam family with a reference protein (for example, a shared protein domain, or a smaller protein being similar to a domain on a longer protein, but failing the 80% bidirectional coverage threshold). 65,507 queries were denoted as *answerable*. Each query protein was also denoted as *remote* if the query had no reference hit above 30% sequence identity and at least 80% query coverage. Overall, 22,076 queries were denoted as both *answerable* and *remote*, forming the denominator for each Test 1 TPR metric in this manuscript.

In order to test the retrieval capabilities of pLMs on negative data, we created sets of synthetic decoys. Specifically, we created 5,000 decoys per class per seed, across 5 different seeds. Each class’ decoys were length-matched to the real query length distribution. The six synthetic classes (D0–D5) are listed in Table 1 alongside the natural hard negatives (D6) described below. By construction, the synthetic decoys are all true negatives with no homologous match against the reference dataset.

**Table 1:** Negative query classes for Test 1. D0–D5 are synthetic decoys; D6 are natural hard negatives.

| Class | Construction | What a hit there means |
|---|---|---|
| D0 | i.i.d. uniform over 20 residues | Score responds to nothing at all |
| D1 | i.i.d. from reference composition | Score responds to bulk composition |
| D2 | Per-sequence shuffle of a real query | Score responds to composition alone, exactly matched |
| D3 | Reversal of a real query | Score responds to composition & reversed local order |
| D4 | Order-2 Markov chain on the reference | Score responds to short *k*-mer statistics |
| D5 | Order-3 Markov chain on the reference | Score responds to slightly longer *k*-mer statistics |
| D6 | Real Swiss-Prot proteins, Pfam clan absent from reference, <30% sequence identity | Score responds to protein-likeness, not homology |

Finally, we also define *natural hard negatives* (D6) as remaining real Swiss-Prot proteins with no detectable relative in the reference. Two criteria had to hold: (i) every Pfam on the protein belongs to a Pfam clan with zero reference members, and (ii) its measured sequence identity to the reference is below 30% (following the previous MMseqs2 clustering and searches). This yielded 3,344 members. We note that these are a weaker ‘true negative’ than synthetic decoys, as a natural hard negative has no *detected* clan-mate. We also note that D6 is shorter than the positives (median length 166 versus 279 residues) and almost entirely single-domain (98.5% versus 75.1% of positives), so a threshold fitted on it may not transfer exactly to the positive population.

#### 2.1.2 Test 2 - SCOPe/ASTRAL-40 Structure Based Splits

We downloaded 15,176 protein domains from SCOPe/ASTRAL v2.08 [Chandonia et al., 2004, 2022], each with expert assessment of protein structure leading to hierarchical SCOPe classifications (Family, Superfamily, Fold and Class levels). We split the dataset holding out at the Family level. 1,176 query families with 4,069 domains were held out as queries, while 3,527 families with 11,107 domains were kept as references. 3,503/4,069 queries were denoted as *answerable* if the query’s Superfamily existed in the reference via another Family (as the same Superfamily should contain protein families with common functional and structural features), forming the denominator for each Test 2 TPR metric in this manuscript. Following SCOPe’s construction, negatives were defined as hits in a different fold, while the ambiguous classification captures SCOPe folds that are *analogous* rather than homologous, as outlined in Table 2.

**Table 2:** Definition of correctness for Test 2 at the individual-hit level.

| **Top hit lands in** | **Verdict** |
|---|---|
| Same superfamily, different family | **Positive** |
| Different fold | **Negative** (defines Test 2’s FPR) |
| Same fold, different superfamily | **Ambiguous** — excluded from both |

Each answerable query’s top hit is labelled first, and a threshold then accepts it if its score is high enough. TPR counts accepted correct-superfamily hits over all 3,503 answerable queries; FPR counts accepted different-fold hits over all different-fold top hits, so its denominator depends on the method. Same-fold, different-superfamily ambiguous hits enter neither rate, and the 566 non-answerable queries are used in neither. Unlike most Test 1 decoys, Test 2’s false positives are real proteins with the wrong fold.

#### 2.1.3 Test 3 - SCOPe/ASTRAL-40 Foldseek Benchmark

Test 3 follows the SCOPe sensitivity benchmark introduced in the Foldseek manuscript [van Kempen et al., 2024], updated for SCOPe v2.08. We chose to include this in Figures S10 and S11 to allow for comparison across the literature as an accepted benchmark in the field, despite the lack of threshold, calibration and decoys that the other tests use.

We used the 13,822/15,176 domains with structures downloadable via `foldseek databases`. Every domain is then searched all-against-all and each ranked hit list is scored up to the first different-fold hit (defined as a false positive). Sensitivity is the fraction of true positives found before that point, where a true positive is a hit from the same family (family level split), same superfamily but different family (superfamily level split), or same fold but different superfamily (fold level split). As in the original benchmark, a domain is scored only if it has a partner at every level, leaving 5,003 query domains. For the embedding arms, the ZCA whitening transform was fit on the 11,107 Test 2 reference domains and applied to all domains; because these domains are also Test 3 queries, this fit is not query-blind.

### 2.2 Method Arms

#### 2.2.1 Sequence-Based Homology Baseline

We used MMseqs2 [Steinegger and Söding, 2017] v18.8cc5c with the parameters -s 7.5 -e 1000 `--max-seqs` 1000 as the representative sequence-based method.

#### 2.2.2 Structure-Based Homology Baseline

We used Foldseek [van Kempen et al., 2024] v10.941cd33 with the parameters `-e` 1000 `-s 9.5 --max-seqs` 1000 as the representative structure-based method using either AlphaFold2 [Jumper et al., 2021] or ASTRAL/PDB [Chandonia et al., 2004, Berman et al., 2000] derived query and reference structures (‘Foldseek’). We also used ProstT5 pLM predicted Foldseek 3Di [Heinzinger et al., 2024] implemented using Phold’s `phold proteins-predict` function as a hybrid pLM-structure method (‘ProstT5-Foldseek’) [Bouras et al., 2026]. For Test 1, we note as a minor limitation that 289,206 / 296,807 (97.44%) of reference proteins had an available Foldseek formatted structure (downloadable via `foldseek databases`), meaning the Foldseek and other methods have slightly different databases, though we expect the impact will be negligible on the results presented.

#### 2.2.3 pLM-based Methods

We tested 10 pLMs: nine general-purpose models (3 across the ESM-2 [Lin et al., 2023] family, 3 across the ESM-C family [ESM Team, 2024], Ankh-base and Ankh-large [Elnaggar et al., 2023] and ProtT5 [Elnaggar et al., 2022]) and the bilingual AA ↔ 3Di language model ProstT5 [Heinzinger et al., 2024]. We also tested gLM2 [Cornman et al., 2025], a genomic language model trained on mixed-modality genomic context and finetuned on retrieval using UniRef50 [Suzek et al., 2007] and the AlphaFold database [Varadi et al., 2024], applied to protein sequences exactly as deployed in Gaia [Jha et al., 2025]. We also tested ProtTucker [Heinzinger et al., 2022], which places a feed-forward neural network atop ProtT5, trained on the CATH [Orengo et al., 1997] hierarchy via contrastive learning. ProtTucker was included as a pseudo-positive control for Test 2, given the similarities between SCOPe and CATH hierarchies used as finetuning data.^3^ All models and checkpoints used can be found in Table S1.

After generating per-residue embeddings for each protein in the query and reference datasets, the final layer was chosen for each model.^4^ Per-residue embeddings were then mean-pooled, yielding a single pooled embedding representation for each protein. For each query, we then retrieved the top 50 nearest neighbour proteins in the reference set with the highest cosine similarity (i.e. top 50 kNN).

### 2.3 Anisotropy

Cosine similarity weights every embedding dimension equally, whereas pLM embeddings may concentrate variance in only a few dimensions. Let *X ∈* ℝ^*n×d*^ denote the *n* mean-pooled reference embeddings of dimension *d*, with mean *µ ∈* ℝ^*d*^ and covariance Σ.

To estimate the number of effective dimensions, we quantified this concentration on each reference database using the *participation ratio* of the covariance eigenspectrum,

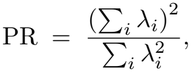

where *λ*_*i*_ is the variance captured along the *i*-th principal direction *v*_*i*_. Note that 1 ≤ PR ≤ *d*, with PR = *d* when the embedding is isotropic.

We then considered four correction methods applied to mean-pooled embeddings to see how they would impact remote homology detection.

- Centering i.e. removing the common mean component but leaving variance and correlation structure unchanged

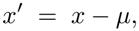
- All-but-the-top (ABTT) [Mu and Viswanath, 2018] i.e. centering followed by projecting out the leading *k* eigenvectors *v*_1_, …, *v*_*k*_ of Σ,

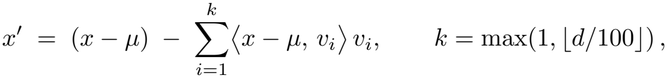

with *k* fixed *a priori* and never tuned, giving e.g. *k* = 6, 7, 10, 15, 25 for *d* = 640, 768, 1024, 1536, 2560. The dominant directions are discarded rather than rescaled.
- Diagonal whitening i.e. centering by mean and rescaling each coordinate by its standard deviation

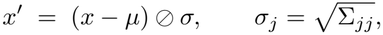

This was used in [Lu et al., 2025] to correct per-channel scale while ignoring inter-channel correlation.
- ZCA whitening [Bell and Sejnowski, 1996] i.e. applying the symmetric whitening transform

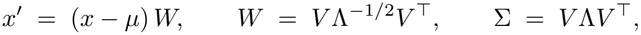

This means the transformed covariance is the identity such that every direction is rescaled to unit variance and none is discarded. It rescales the dominant dimensions rather than discarding them like ABTT, and cosine similarity after whitening equals cosine similarity under the Mahalanobis inner product of the reference set. We used exact ZCA without shrinkage, flooring eigenvalues at 10^*−*8^ before inversion; shrinking the covariance towards a scaled identity changed Test 2 sensitivity by −8.9 to +9.0 points depending on the model, with no consistent direction (Table S4).

All four transforms are estimated on only the reference database in each analysis.^5^

## 3 Results

### 3.1 pLM Cosine Similarity Struggles to Differentiate True Positives From Negatives

Our first analysis was to determine the sensitivity of retrieval for all methods holding at a 1% FPR across the natural hard negatives (Test 1) and wrong-fold hits (Test 2), presented in Figure 1. Using

**Figure 1:**
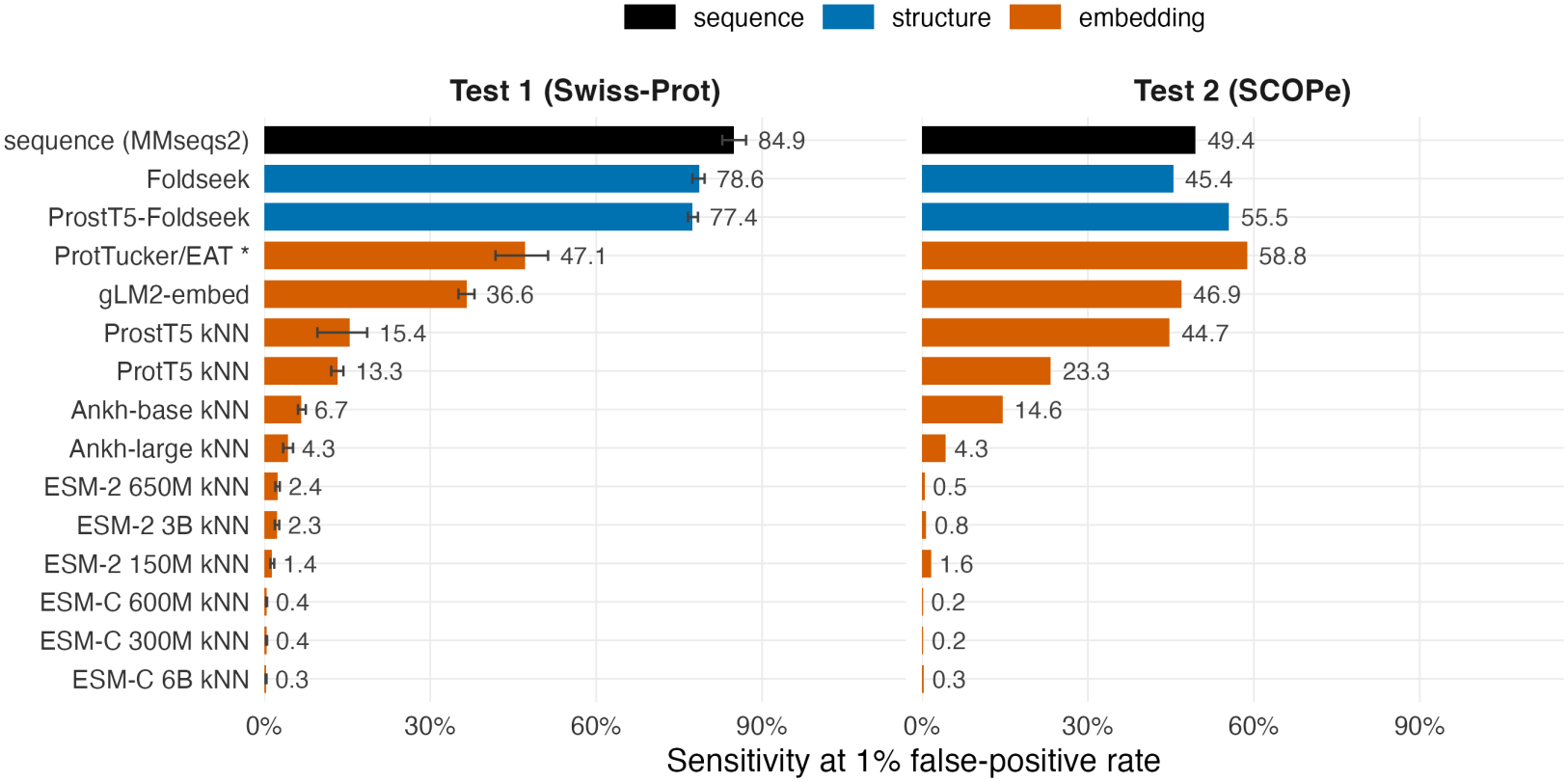
Sensitivity of retrieving true positive hits at 1% false-positive rate for all methods. Test 1 bars are means over 400 query-level bootstrap replicates with the threshold refitted in each; whiskers are 95% percentile intervals. Test 2 uses a single threshold at 1% different-fold FPR; sensitivity is over all 3,503 answerable queries. Note that Test 1 was curated with MMseqs2, so its sequence result is an upper bound. *\** ProtTucker is contrastively trained on CATH, which classifies the same structures as SCOPe.

Test 1 as an example, we set the threshold at the score (cosine similarity, or E-value) of the 34th-highest-scoring of the 3,344 natural hard negatives against its closest reference hit, so that 1.02% of negatives pass. A sensitivity of 50% means that 50% of the 22,076 answerable and remote queries had their closest hit at or above this threshold.

On Test 1, all embedding-based models performed worse than sequence and structure-based methods (Figure 1), with MMseqs2 unsurprisingly performing the best (as the dataset was constructed using it). ProstT5-Foldseek substantially outperformed ProstT5 kNN, while the two finetuned methods gLM2-embed and ProtTucker performed more strongly than the other methods. ESM-C across all sizes performed the worst. Test 2 showed a similar result, though gLM2-embed and ProtTucker performed comparably with structure and sequence based methods, and ProstT5 kNN performed much better on this structure-based dataset. Test 2 orderings depend on the FPR convention: counting false positives against all 3,503 answerable queries rather than against each method’s different-fold top hits (as in Figure 1), ProstT5 kNN (57.8%) and gLM2-embed (57.2%) exceed MMseqs2 (50.6%). ProstT5-Foldseek outperformed Foldseek on Test 2 (55.5 versus 45.4%), also when restricted to queries both searched (56.6 versus 49.4%), possibly because experimental ASTRAL coordinates are incomplete domain fragments while predicted 3Di covers every residue.

These conclusions are robust to how thresholds and positives are defined. On the 566 non-answerable Test 2 queries, which cannot have a correct hit, the Figure 1 thresholds give 1.1% false positives for MMseqs2 and Foldseek but 3.8–4.7% for raw ESM-C (Table S6). On Test 1, restricting positives to single-domain queries, or to queries whose set of Pfam families occurs in the reference, changes no method by more than 6.1 points (Table S7). Reweighting D6 to the positives’ length distribution leaves MMseqs2 unchanged (83.9%) but lowers ProtTucker to 30.4% and ProstT5 kNN to 6.3%, suggesting the length mismatch between calibration negatives and queries moves embedding thresholds but not E-value thresholds. Finally, 17 of the 34 D6 proteins accepted by MMseqs2 have significant Foldseek hits (base rate 6.7%), against at most 3 for most pure pLMs (Table S7), so some D6 ‘false positives’ are undetected homologs and the alignment advantage is, if anything, understated.

We next looked at the densities of cosine similarities and E-values of the different query classes across different methods. Sequence- and structure-based methods with null models (MMseqs2 and ProstT5-Foldseek) clearly separate remote homologs from negative classes (Figure 2). gLM2-embed was also able to generally distinguish the different classes, with Ankh-base to a lesser degree. For ESM-2 and ESM-C, the D2 shuffled decoy queries generally have higher cosine similarities than real remote homologs. Similar results were observed across all models (Figure S1), all decoy subsets in Test 1 (Figure S2) and also on Test 2 (Figure S3).

**Figure 2:**
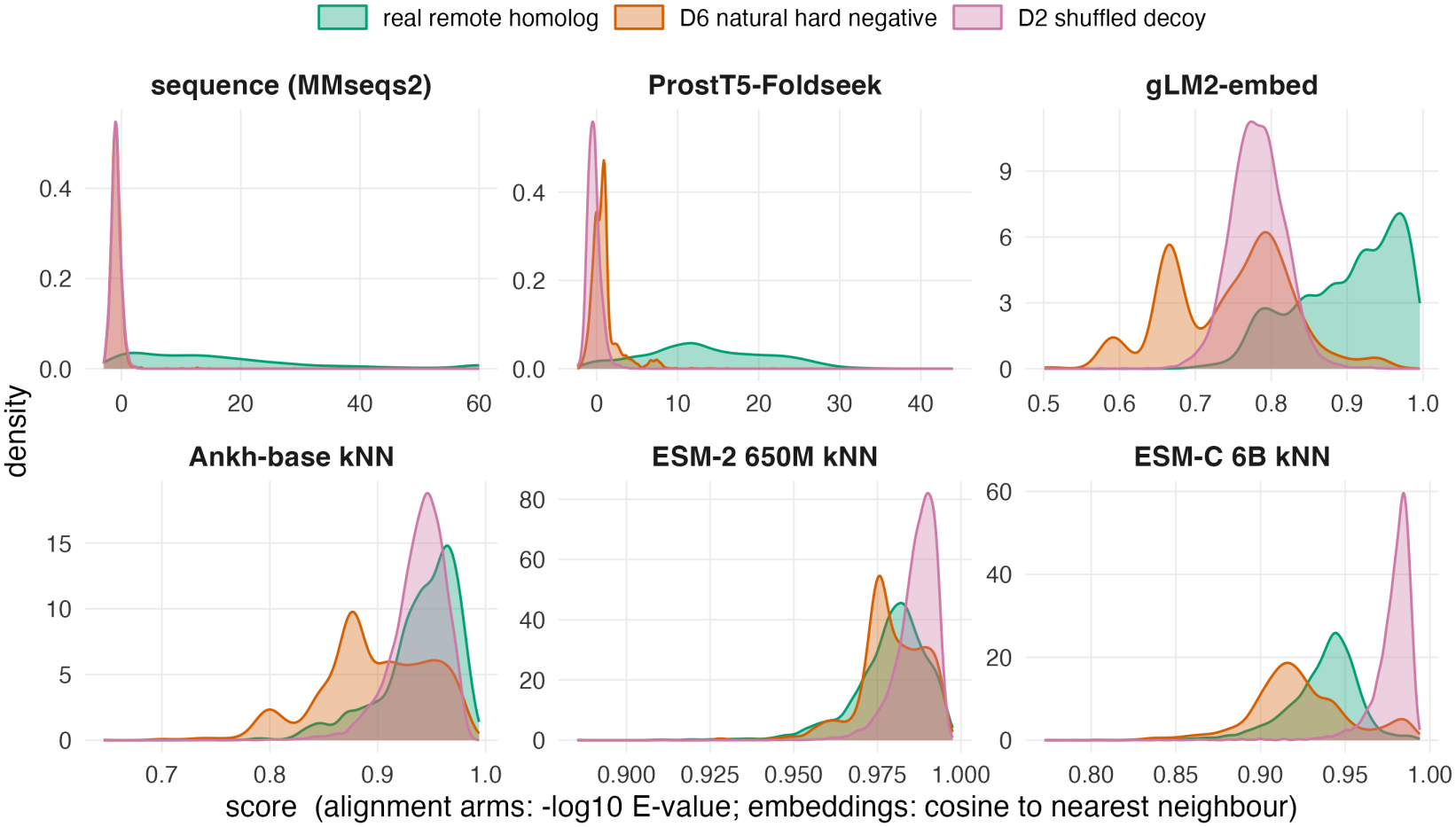
Density of best-hit scores on Test 1 for real remote homologs, shuffled decoys (D2) and natural hard negatives (D6), for selected methods. Alignment arms show −log_10_ E-value capped at 60 for display; embedding arms show cosine similarity to the nearest neighbour. Axes are per panel because the scales are not comparable.

We then looked at the performance of each approach on queries stratified by length. Before conducting the analysis, we had hypothesised that short proteins (below 100 and certainly below 50 AA) would have a much higher false positive rate and lower sensitivity, due to the limits of pLMs [Ding, 2026] and structure prediction methods in representing short proteins and peptides [Monzon et al., 2022]. All methods have a low false positive rate below 100 AA other than ProtT5, ProstT5 and ProtTucker, and ProtT5 also had a high sensitivity on the shortest 1–49 AA bin (Figure S4). The structure-based methods had low sensitivity <100AA (under 10%), but matched MMseqs2 in the largest length bin. Above 50 AA, ESM-2 650M and ESM-C 6B exhibited near 0% sensitivity.

We next computed the fraction of the reference set ever retrieved (i.e. the proportion of all reference proteins that appear among the 50 nearest neighbours across the set of query proteins), for real remote homology and decoy query sets. We hypothesised for strongly performing models that real remote homologs and decoys would have a similar spread of reference protein hits among the top 50 hits, while decoys for poorly performing models would also hit the same references likely to be found in the ‘embedding junkyard’ of low quality embeddings [Prabakaran and Bromberg, 2026].

For Test 2, real queries reach 82–98% of the reference database in every model, whereas the decoys span 5.3–64.7% for every model except ProtTucker (95.3%) (Figure S5). Results for Test 1 are similar. gLM2-embed has much more similar coverage between real and decoy queries, whereas for ESM-C 6B, a single 97-residue SCOPe domain (d6z4ua_, b.164.1.1) sits in the top-50 of 70.4% of ESM-C 6B’s 20,345 shuffled peptides, against 0.6% of its real queries. This suggests that decoys are drawn into ESM-C 6B’s ‘embedding junkyard’ with high cosine similarities, implying that if you don’t know if your protein has a good quality embedding *a priori*^6^, retrieval searches may produce many high cosine similarity but spurious highly ranked hits.

### 3.2 Correcting for Anisotropy Improves pLM Remote Homology Detection

Considering the poor performance of pure pLMs on our benchmarks and datasets, we then assessed the effect of embedding-space anisotropy on the inflated cosine similarities on decoy proteins compared to real homologs. Anisotropy is well documented in contextual language-model embeddings [Ethayarajh, 2019], and whitening mean-pooled sentence embeddings is an established fix for cosine retrieval in NLP [Su et al., 2021]. While this transform is standard, it has received limited attention in the pLM literature, particularly regarding its effect on calibrated retrieval of protein homologs. Hakim et al. [2025] reported ProtBERT is highly anisotropic. Lu et al. [2025] showed that pLMs in the ESM-2 family show massive activations in some dimensions, while McWhite et al. [2023] showed using batch correction methods could improve detection of reciprocal best hits across retrieval tasks.

We first calculated the participation ratio to measure the number of effective dimensions. On Test 2’s dataset, ESM-C models occupied only PR = 3.3–5.1 of a nominal *d* = 960–2560 (0.20–0.34%) and ESM-2 650M PR = 6.0 of *d* = 1280 (0.47%), an order of magnitude less than ProtT5, which occupied 63.0 of 1024 (6.2%) and gLM2-embed 30.6 of 512 (6.0%). Additionally, the top dimension contributes 54.3% of variance for ESM-C 300M, whereas for ProtT5, this was only 7.0%. See Tables S2 and S3 for all models. Across the eleven models other than ProtTucker, PR is strongly associated with Test 2 sensitivity at 1% FPR (Spearman *ρ* = 0.89), though we note that Ankh-large has the second-highest PR of the eleven yet gains the most from whitening. Interestingly, removing the shared mean rarely helps (centering was worse than baseline for 7 of 11 models on Test 2), and for the models we measured (ESM-2 650M and ESM-C 6B), shuffled decoys lie further from the reference centroid than real queries, so a common offset is not the problem. Nor are the dominant directions length or composition, which explain 7–14% of first-PC variance in ESM-2 650M and ESM-C; regressing them out does not restore real-versus-shuffle separation (Table S8).

We then considered the impact of four correction methods (centering, ABTT, diagonal whitening and ZCA whitening) on the benchmark outlined in Figure 1. On Test 2, ZCA whitening improved the sensitivity of all 10 pure pLM methods and gLM2-embed, and was the best-performing transform for 9 of 11 models, with ABTT performing best on the other two (Ankh-base and ESM-2 3B) (Figure S6). The largest improvements were on Ankh and ESM-2 family models. Similar improvements were shown on Test 1 also (Figure S7). After whitening, real homologs outrank synthetic decoys on Test 1 for eight of eleven models (median AUROC over synthetic classes from 0.74 for ESM-2 650M to 0.89 for Ankh-base), but ESM-C remains below chance (0.29–0.35). Nor does one threshold hold across negative types: at the 1% threshold fitted on natural negatives, raw pLMs admit up to 27% of a synthetic decoy class (ESM-C 6B, order-2 Markov) against at most 0.7% for alignment-based and retrieval-trained methods, and whitening redistributes rather than removes this leakage (Table S5).

We next looked at the impact of ZCA whitening on the more permissive Top-1 retrieval analysis on Test 2 (i.e. the percentage of queries whose top-ranked hit shares the query’s superfamily, with no threshold applied at all to potentially control false positives) (Figure 3). ZCA whitening markedly improved Top-1 retrieval across most models, particularly the ESM-2 and ESM-C families and ProtT5. After whitening, every model other than the ESM-C family and gLM2-embed reached a Top-1 rate between 85.3 and 88.8%, comparable to the unwhitened Foldseek (80.1%) and ProtTucker (89.0%). As expected from a sequence-alignment-only method, MMseqs2 had lower Top-1 retrieval (56.9%) than every whitened embedding model. Figures S8 and S9 show the impact of all four transforms on the Top-1 metric across both Test 1 and Test 2, with ZCA whitening performing best for every model except gLM2-embed, where ABTT is marginally higher, and showing strong improvements over baseline.

**Figure 3:**
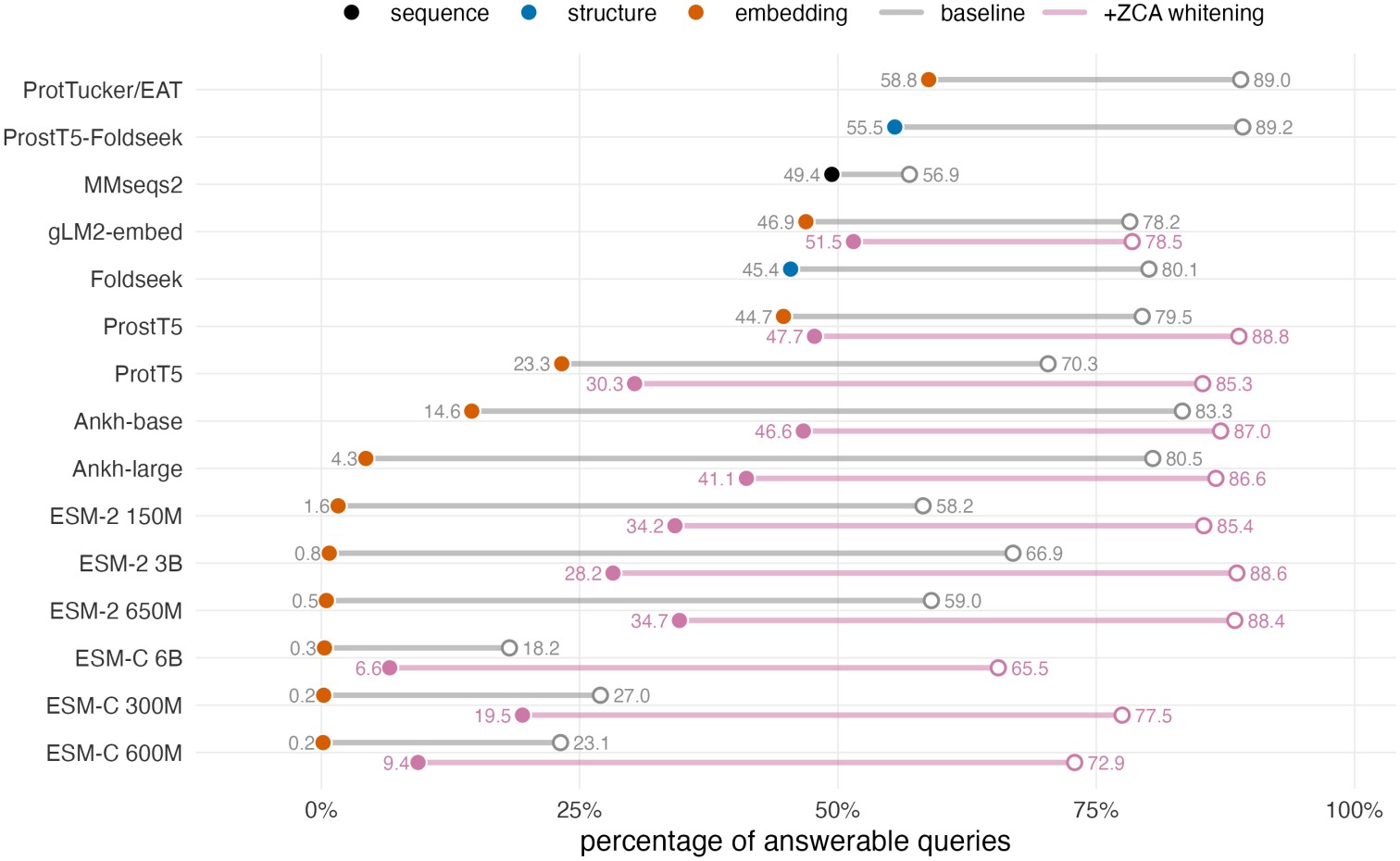
Test 2 (SCOPe) sensitivity at 1% FPR (filled dots) and Top-1 retrieval (hollow dots) across all methods. Threshold-free Top-1 retrieval shows a higher retrieval performance on all models compared to the thresholded sensitivity metric. For pLM methods, both metrics after ZCA whitening are shown in pink and the unwhitened models in grey.

Finally, on the widely used all-against-all Foldseek SCOPe benchmark i.e. Test 3, ZCA whitening also improves all pure pLM methods at the family and superfamily levels, with only gLM2-embed performing worse after whitening, though at the fold level it helps little (Figures S10 and S11). While not reaching the sensitivity of Foldseek across the different Family, Superfamily and Fold levels, this shows that correcting for anisotropy is a simple and fast way to improve remote homology detection on well established benchmarks in the field.

## 4 Conclusion and Limitations

We have shown in this work that it is difficult to generate reasonable cosine similarity thresholds to conduct nearest neighbour retrieval using mean-pooled embeddings (as commonly conducted in the field) that control potential false positives. No single cosine threshold separates true from false hits at a controlled false-positive rate, and for the widely used ESM-2 and ESM-C families, shuffled sequences outscore real remote homologs. Anisotropy and specifically low effective dimensionality is strongly associated with this failure, and whitening the embeddings against the reference database is a cheap correction that substantially improves ranking. However, it does not supply what alignment tools have, i.e. a null model that makes scores comparable across queries.

While the curation of our test datasets (Test 1 based on Swiss-Prot and Test 2 based on SCOPe) may not be perfect, considering the difficulty of creating benchmarks in the remote homology detection and protein annotation fields^7^, we think nonetheless they show the impact of anisotropy on realistic embedding-based annotation. Specifically, Top-1 on Test 1 (lookup against Swiss-Prot) above trustworthy thresholds using sequence homology-based alignment tools is more or less how protein annotation is commonly conducted in practice in widely used software packages (e.g. Prokka [Seemann, 2014] and Bakta [Schwengers et al., 2021]).

Our conclusions apply to final-layer, mean-pooled embeddings compared by cosine similarity. Other layers and pooling schemes [NaderiAlizadeh and Singh, 2025], local embedding alignment methods such as pLM-BLAST [Kaminski et al., 2023], and retrieval-trained models such as TM-Vec [Hamamsy et al., 2024], PLMSearch [Liu et al., 2024] and LEMON-ZEST [Banerjee et al., 2026] may behave differently and should be benchmarked the same way. We also did not test temporally held-out negatives, or functional endpoints such as GO or EC transfer. Finally, Test 1 could be expanded to proteins longer than 400 AA.

## Acknowledgments and Disclosure of Funding

This work was supported with the assistance of resources and services from Phoenix HPC at Adelaide University and Pawsey Supercomputing Research Centre, which is supported by the Australian Government. We would like to thank Fabien Voisin and Sarah Beecroft for their assistance at Phoenix and Pawsey, respectively.

## LLM Use

We declare that LLMs (specifically, Claude Opus 5 and Fable 5) were used to help orchestrate analysis presented in this manuscript (specifically, most of the code for running analyses and making the first draft of figures). It also was used for ideation in the anisotropy analysis. However, the entire manuscript was written by us humans, with some light LLM assistance for editing, proofreading and format compliance.

## Reproducibility

All code is available at https://github.com/gbouras13/know-thy-neighbour, and intermediate data and embeddings are available on Zenodo at https://doi.org/10.5281/zenodo.24181855.

## A Supplementary Tables

**Table S1:**
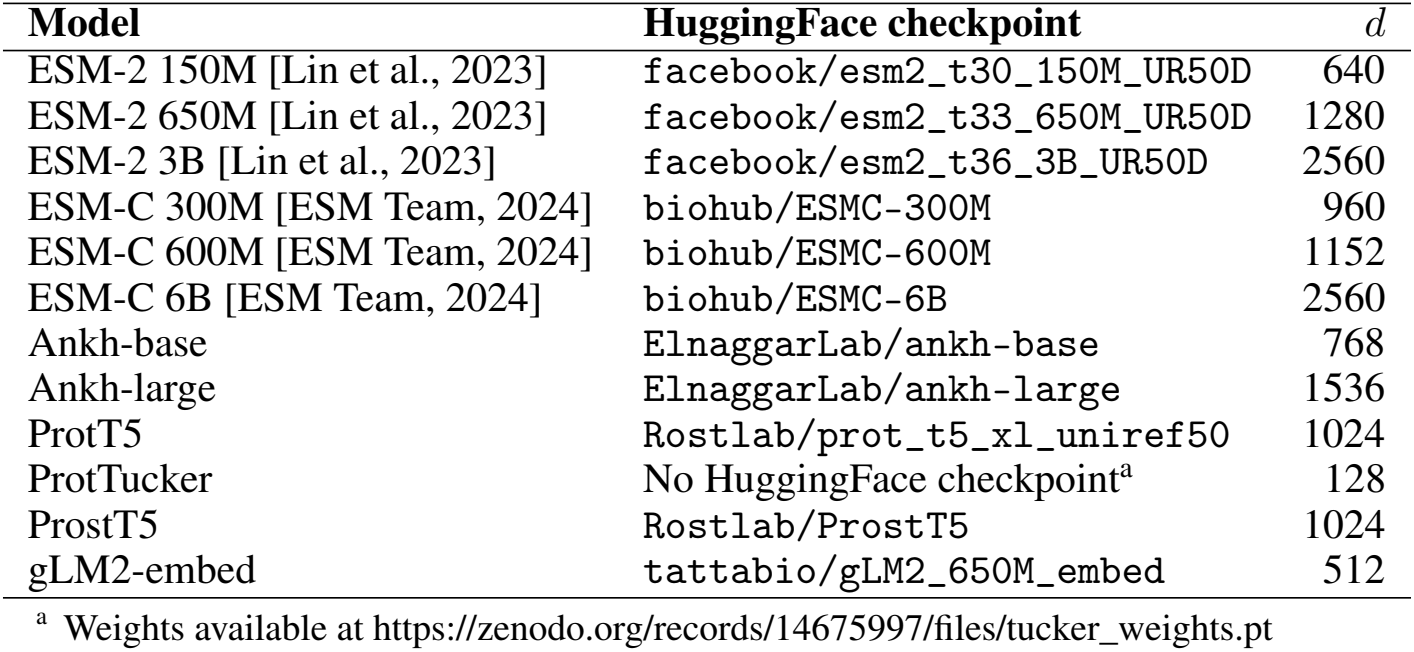
pLM variants and embedding dimensions used in this study. All models were run in bfloat16 except ESM-2 variants, which were run in float16. Note ProtTucker is a supervised 128-d FFN projection atop ProtT5, not a raw pLM output.

**Table S2:** Anisotropy on the Test 1 (Swiss-Prot) reference set, *n* = 296,807. ^*†*^ProtTucker is a supervised 128-d projection of ProtT5, not a raw pLM output.

|  |  | effective dim. |  | *k* for var. |  | top-1 |
|---|---|---|---|---|---|---|
| Model | *d* | PR | % of *d* | 50% | 90% | var. % |
| ProtTucker/EAT<sup>†</sup> | 128 | 56.1 | 43.81 | 30 | 97 | 8.4 |
| Ankh-large | 1536 | 52.4 | 3.41 | 37 | 431 | 8.4 |
| ProtT5 | 1024 | 50.9 | 4.97 | 24 | 247 | 8.4 |
| Ankh-base | 768 | 39.8 | 5.18 | 24 | 206 | 10.3 |
| gLM2-embed | 512 | 33.9 | 6.61 | 23 | 137 | 13.0 |
| ProstT5 | 1024 | 18.9 | 1.85 | 8 | 160 | 17.2 |
| ESM-2 3B | 2560 | 14.1 | 0.55 | 12 | 388 | 25.0 |
| ESM-C 6B | 2560 | 10.9 | 0.43 | 5 | 119 | 26.5 |
| ESM-2 150M | 640 | 10.8 | 1.69 | 6 | 78 | 27.4 |
| ESM-2 650M | 1280 | 8.3 | 0.64 | 5 | 146 | 32.5 |
| ESM-C 600M | 1152 | 5.6 | 0.48 | 3 | 54 | 40.3 |
| ESM-C 300M | 960 | 4.8 | 0.50 | 2 | 50 | 44.2 |

**Table S3:** Anisotropy on the Test 2 (SCOPe/ASTRAL-40 2.08) reference set, *n* = 11,107. ^*†*^ProtTucker is a supervised 128-d projection of ProtT5, not a raw pLM output. Notably, PR is strongly correlated with Test 2 sensitivity at 1% FPR (Figure 1), Spearman’s *ρ* = +0.891. Ankh-large is the primary counterexample, with the second-highest PR of these eleven (44.8) but with low sensitivity (see Figure 1).

|  |  | effective dim. |  | *k* for var. |  | top-1 |
|---|---|---|---|---|---|---|
| Model | *d* | PR | % of *d* | 50% | 90% | var. % |
| ProtT5 | 1024 | 63.0 | 6.16 | 29 | 277 | 7.0 |
| ProtTucker/EAT<sup>†</sup> | 128 | 56.0 | 43.76 | 34 | 101 | 8.0 |
| Ankh-large | 1536 | 44.8 | 2.92 | 43 | 516 | 10.2 |
| gLM2-embed | 512 | 30.6 | 5.98 | 27 | 161 | 14.5 |
| ProstT5 | 1024 | 27.1 | 2.65 | 11 | 214 | 11.5 |
| Ankh-base | 768 | 26.8 | 3.49 | 23 | 228 | 14.8 |
| ESM-2 150M | 640 | 9.9 | 1.55 | 6 | 88 | 29.3 |
| ESM-2 3B | 2560 | 8.6 | 0.34 | 8 | 482 | 33.3 |
| ESM-2 650M | 1280 | 6.0 | 0.47 | 3 | 161 | 39.1 |
| ESM-C 6B | 2560 | 5.1 | 0.20 | 2 | 89 | 42.9 |
| ESM-C 600M | 1152 | 3.7 | 0.32 | 1 | 45 | 51.1 |
| ESM-C 300M | 960 | 3.3 | 0.34 | 1 | 44 | 54.3 |

**Table S4:** Test 2 sensitivity (%) at 1% FPR under exact ZCA, as used throughout, and with linear shrinkage of the reference covariance towards a scaled identity, 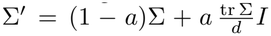 with *a* = *d/n* (up to 0.23 for *d* = 2560, *n* = 11,107). Eigenvalues are floored at 10^*−*8^ in both.

| Model | exact ZCA | shrunk ZCA | Δ |
|---|---|---|---|
| ESM-2 3B | 28.2 | 37.2 | +9.0 |
| ESM-2 650M | 34.7 | 39.9 | +5.2 |
| Ankh-large | 41.1 | 45.7 | +4.6 |
| Ankh-base | 46.6 | 49.7 | +3.1 |
| ProstT5 | 47.7 | 50.4 | +2.7 |
| gLM2-embed | 51.5 | 51.7 | +0.2 |
| ESM-2 150M | 34.2 | 32.1 | −2.1 |
| ProtT5 | 30.3 | 27.0 | −3.3 |
| ESM-C 600M | 9.4 | 4.5 | −4.9 |
| ESM-C 6B | 6.6 | 1.5 | −5.1 |
| ESM-C 300M | 19.5 | 10.6 | −8.9 |

**Table S5:** Achieved false-positive rate (%) on each Test 1 negative class at the 1% threshold fitted on the natural hard negatives (D6), as in Figure 1; bootstrap means over 400 replicates, all length bins pooled. Classes are defined in Table 1. The D6 column is the calibration target at 1% (and so is just over 1%).

| Method | D0 | D1 | D2 | D3 | D4 | D5 | D6 |
|---|---|---|---|---|---|---|---|
| MMseqs2 | 0.39 | 0.52 | 0.38 | 0.68 | 0.52 | 0.54 | 1.04 |
| Foldseek | 0.00 | 0.00 | 0.00 | 0.00 | 0.00 | 0.00 | 1.03 |
| ProstT5-Foldseek | 0.00 | 0.00 | 0.00 | 0.01 | 0.00 | 0.00 | 1.03 |
| gLM2-embed | 0.00 | 0.00 | 0.04 | 0.14 | 0.00 | 0.00 | 1.03 |
| ProtTucker | 0.00 | 0.00 | 0.00 | 0.02 | 0.00 | 0.00 | 1.03 |
| ProtT5 | 0.06 | 0.03 | 0.32 | 0.41 | 0.24 | 0.21 | 1.03 |
| ProstT5 | 0.00 | 0.24 | 0.12 | 0.20 | 0.43 | 0.39 | 1.03 |
| Ankh-base | 0.05 | 1.25 | 0.81 | 2.53 | 2.92 | 3.02 | 1.04 |
| Ankh-large | 3.48 | 8.21 | 6.02 | 10.78 | 10.06 | 10.37 | 1.03 |
| ESM-2 150M | 0.01 | 0.55 | 0.22 | 1.08 | 0.82 | 0.98 | 1.03 |
| ESM-2 650M | 0.26 | 3.02 | 1.21 | 2.74 | 4.32 | 5.01 | 1.04 |
| ESM-2 3B | 0.15 | 3.22 | 1.38 | 3.66 | 4.89 | 5.69 | 1.03 |
| ESM-C 300M | 0.54 | 4.95 | 2.17 | 3.35 | 11.10 | 10.95 | 1.03 |
| ESM-C 600M | 0.09 | 3.34 | 1.40 | 3.00 | 7.51 | 7.89 | 1.03 |
| ESM-C 6B | 1.64 | 13.97 | 5.93 | 6.86 | 27.11 | 25.34 | 1.04 |
| gLM2-embed +whiten | 0.00 | 0.01 | 0.02 | 0.05 | 0.01 | 0.01 | 1.04 |
| ProtT5 +whiten | 0.28 | 0.24 | 0.40 | 0.29 | 0.74 | 0.52 | 1.03 |
| ProstT5 +whiten | 0.16 | 0.13 | 0.37 | 0.90 | 0.20 | 0.15 | 1.04 |
| Ankh-base +whiten | 2.22 | 0.84 | 1.05 | 0.26 | 0.85 | 0.66 | 1.03 |
| Ankh-large +whiten | 0.68 | 2.16 | 2.91 | 0.76 | 2.01 | 1.38 | 1.03 |
| ESM-2 150M +whiten | 0.14 | 1.13 | 1.71 | 0.64 | 1.79 | 1.26 | 1.03 |
| ESM-2 650M +whiten | 0.25 | 2.89 | 3.75 | 1.21 | 2.89 | 2.09 | 1.04 |
| ESM-2 3B +whiten | 0.21 | 0.81 | 1.10 | 0.28 | 1.11 | 0.82 | 1.03 |
| ESM-C 300M +whiten | 0.96 | 7.92 | 7.63 | 1.82 | 3.65 | 2.66 | 1.03 |
| ESM-C 600M +whiten | 6.81 | 11.03 | 8.37 | 1.91 | 5.53 | 4.02 | 1.04 |
| ESM-C 6B +whiten | 20.01 | 5.97 | 6.04 | 2.10 | 5.11 | 3.93 | 1.03 |

**Table S6:**
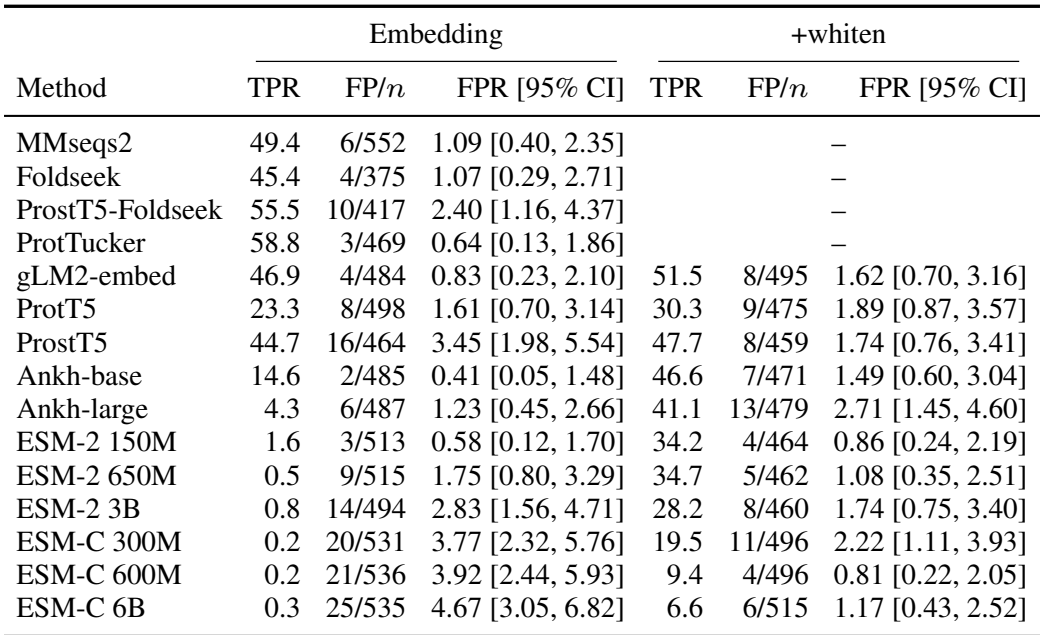
Test 2 thresholds applied to independent negatives. Each method’s 1% FPR threshold is fitted on the 3,503 answerable queries exactly as in Figure 1 and applied unchanged to the 566 non-answerable queries. These are queries whose superfamily is absent from the reference, so any accepted different-fold top hit is a false positive. Same-fold top hits and queries without a hit are excluded, so *n* differs by method.

| Method | Embedding |  |  | +whiten |  |  |
|---|---|---|---|---|---|---|
|  | TPR | FP/*n* | FPR [95% CI] | TPR | FP/*n* | FPR [95% CI] |
| MMseqs2 | 49.4 | 6/552 | 1.09 [0.40, 2.35] | – |  |  |
| Foldseek | 45.4 | 4/375 | 1.07 [0.29, 2.71] | – |  |  |
| ProstT5-Foldseek | 55.5 | 10/417 | 2.40 [1.16, 4.37] | – |  |  |
| ProtTucker | 58.8 | 3/469 | 0.64 [0.13, 1.86] | – |  |  |
| gLM2-embed | 46.9 | 4/484 | 0.83 [0.23, 2.10] | 51.5 | 8/495 | 1.62 [0.70, 3.16] |
| ProtT5 | 23.3 | 8/498 | 1.61 [0.70, 3.14] | 30.3 | 9/475 | 1.89 [0.87, 3.57] |
| ProstT5 | 44.7 | 16/464 | 3.45 [1.98, 5.54] | 47.7 | 8/459 | 1.74 [0.76, 3.41] |
| Ankh-base | 14.6 | 2/485 | 0.41 [0.05, 1.48] | 46.6 | 7/471 | 1.49 [0.60, 3.04] |
| Ankh-large | 4.3 | 6/487 | 1.23 [0.45, 2.66] | 41.1 | 13/479 | 2.71 [1.45, 4.60] |
| ESM-2 150M | 1.6 | 3/513 | 0.58 [0.12, 1.70] | 34.2 | 4/464 | 0.86 [0.24, 2.19] |
| ESM-2 650M | 0.5 | 9/515 | 1.75 [0.80, 3.29] | 34.7 | 5/462 | 1.08 [0.35, 2.51] |
| ESM-2 3B | 0.8 | 14/494 | 2.83 [1.56, 4.71] | 28.2 | 8/460 | 1.74 [0.75, 3.40] |
| ESM-C 300M | 0.2 | 20/531 | 3.77 [2.32, 5.76] | 19.5 | 11/496 | 2.22 [1.11, 3.93] |
| ESM-C 600M | 0.2 | 21/536 | 3.92 [2.44, 5.93] | 9.4 | 4/496 | 0.81 [0.22, 2.05] |
| ESM-C 6B | 0.3 | 25/535 | 4.67 [3.05, 6.82] | 6.6 | 6/515 | 1.17 [0.43, 2.52] |

**Table S7:**
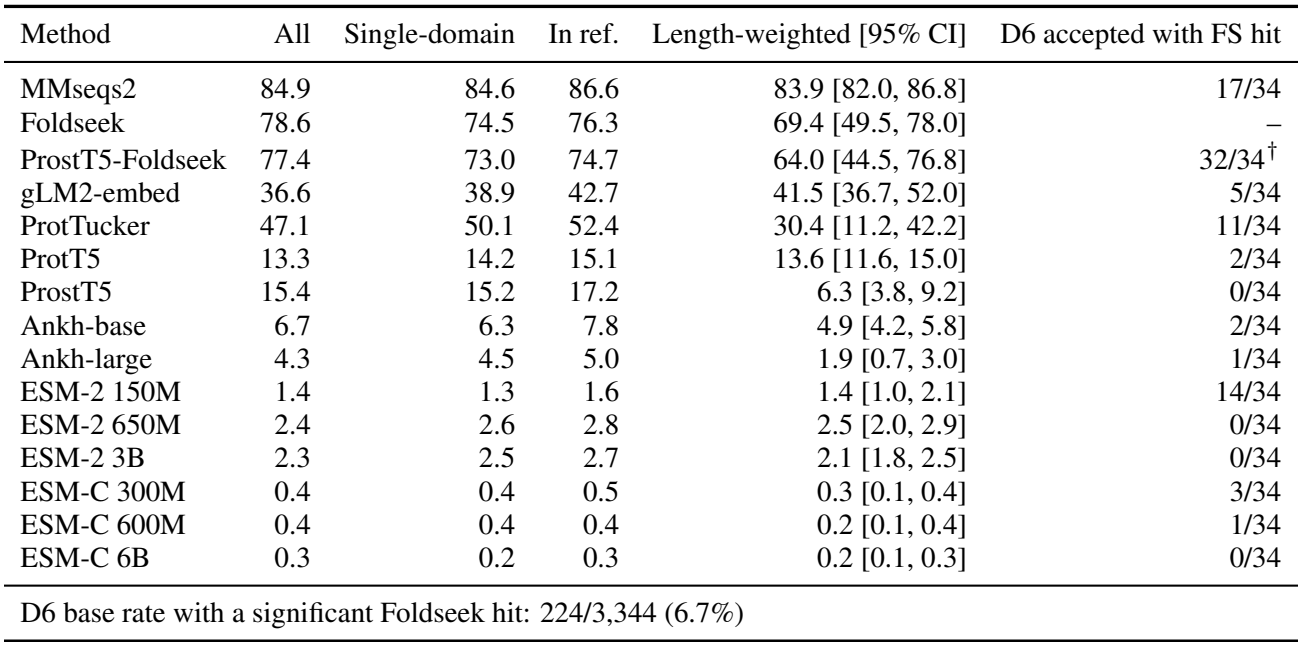
Robustness of Test 1 sensitivity (%) at 1% FPR. *All*: as in Figure 1 (22,076 positives). *Single-domain*: positives with exactly one Pfam family (16,581). *In ref*.: positives whose set of Pfam families also occurs in at least one reference protein (17,797). *Length-weighted*: D6 reweighted per length bin to the positives’ length distribution before fitting the threshold; the weighted 1% rests on few effective negatives. Bootstrap means over 400 replicates. Every method remains significantly below MMseqs2 in every column. *Last column*: of the 34 D6 proteins each method’s threshold accepts, the number with a Foldseek hit at *E <* 10^*−*3^ to the reference; ^*†*^ProstT5-Foldseek is somewhat circular, as the method is uses a Foldseek search; Foldseek itself is not assessed for the same reason.

| Method | All | Single-domain | In ref. | Length-weighted [95% CI] | D6 accepted with FS hit |
|---|---|---|---|---|---|
| MMseqs2 | 84.9 | 84.6 | 86.6 | 83.9 [82.0, 86.8] | 17/34 |
| Foldseek | 78.6 | 74.5 | 76.3 | 69.4 [49.5, 78.0] | – |
| ProstT5-Foldseek | 77.4 | 73.0 | 74.7 | 64.0 [44.5, 76.8] | 32/34<sup>†</sup> |
| gLM2-embed | 36.6 | 38.9 | 42.7 | 41.5 [36.7, 52.0] | 5/34 |
| ProtTucker | 47.1 | 50.1 | 52.4 | 30.4 [11.2, 42.2] | 11/34 |
| ProtT5 | 13.3 | 14.2 | 15.1 | 13.6 [11.6, 15.0] | 2/34 |
| ProstT5 | 15.4 | 15.2 | 17.2 | 6.3 [3.8, 9.2] | 0/34 |
| Ankh-base | 6.7 | 6.3 | 7.8 | 4.9 [4.2, 5.8] | 2/34 |
| Ankh-large | 4.3 | 4.5 | 5.0 | 1.9 [0.7, 3.0] | 1/34 |
| ESM-2 150M | 1.4 | 1.3 | 1.6 | 1.4 [1.0, 2.1] | 14/34 |
| ESM-2 650M | 2.4 | 2.6 | 2.8 | 2.5 [2.0, 2.9] | 0/34 |
| ESM-2 3B | 2.3 | 2.5 | 2.7 | 2.1 [1.8, 2.5] | 0/34 |
| ESM-C 300M | 0.4 | 0.4 | 0.5 | 0.3 [0.1, 0.4] | 3/34 |
| ESM-C 600M | 0.4 | 0.4 | 0.4 | 0.2 [0.1, 0.4] | 1/34 |
| ESM-C 6B | 0.3 | 0.2 | 0.3 | 0.2 [0.1, 0.3] | 0/34 |
| D6 base rate with a significant Foldseek hit: 224/3,344 (6.7%) |  |  |  |  |  |

**Table S8:** What the dominant directions encode (Test 2 reference embeddings). Length + comp. is log length, GRAVY hydrophobicity and the 20 amino-acid fractions. *Regressed*: these covariates (with an intercept, so also centred) regressed out of every embedding, fitted on the reference; AUROC is for real queries against their shuffles (D2); below 0.5, shuffles score higher than real queries.

|  | PC1 share | *R*<sup>2</sup> of PC1 on | Total variance | AUROC real vs. shuffled |  |
|---|---|---|---|---|---|
| Model | of variance | length + comp. | length + comp. | raw | regressed |
| ESM-2 650M | 39.1% | 0.09 | 15.1% | 0.174 | 0.030 |
| ESM-C 300M | 54.3% | 0.14 | 20.9% | 0.035 | 0.016 |
| ESM-C 6B | 43.0% | 0.07 | 17.3% | 0.020 | 0.010 |
| Ankh-large | 10.2% | 0.04 | 13.2% | 0.252 | 0.152 |
| ProtT5 | 7.0% | 0.17 | 16.2% | 0.620 | 0.335 |
| ProstT5 | 11.5% | 0.26 | 20.3% | 0.608 | 0.170 |
| gLM2-embed | 14.5% | 0.60 | 17.3% | 0.739 | 0.528 |

## B Supplementary Figures

**Figure S1:**
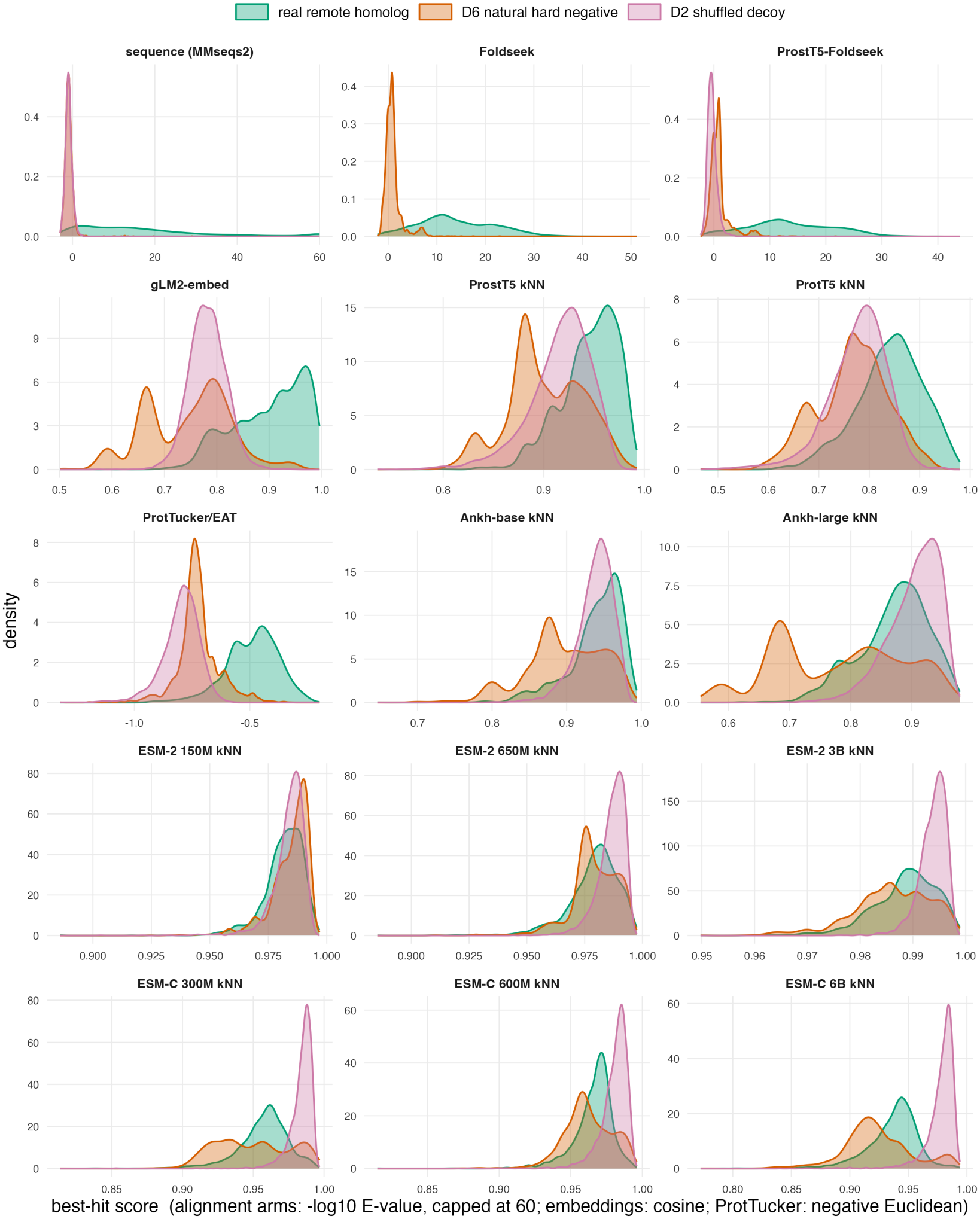
Density plots of cosine similarities/E-values of all methods on Test 1 for queries with real remote homologs, shuffled decoys, and natural hard negatives. Note ProtTucker uses Euclidean distance, stored negated so higher-is-better uniformly. Note also that Foldseek cannot have shuffled queries (as it requires query structures).

**Figure S2:**
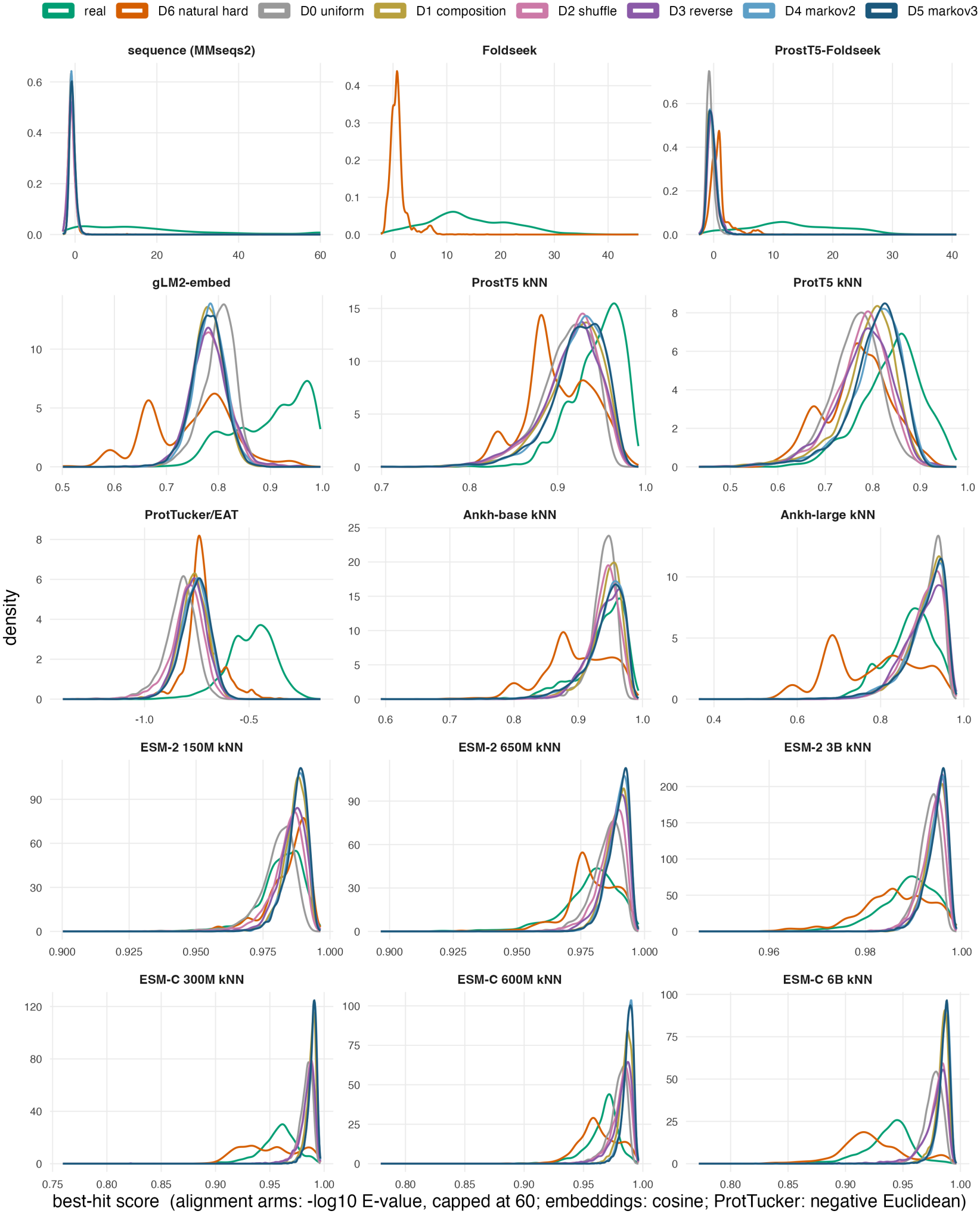
Density plots of cosine similarities/E-values of all methods on Test 1 for all classes of queries with real remote homologs, shuffled decoys, and natural hard negatives. Note ProtTucker uses Euclidean distance, stored negated so higher-is-better uniformly. Note also that Foldseek cannot have shuffled queries (as it requires query structures).

**Figure S3:**
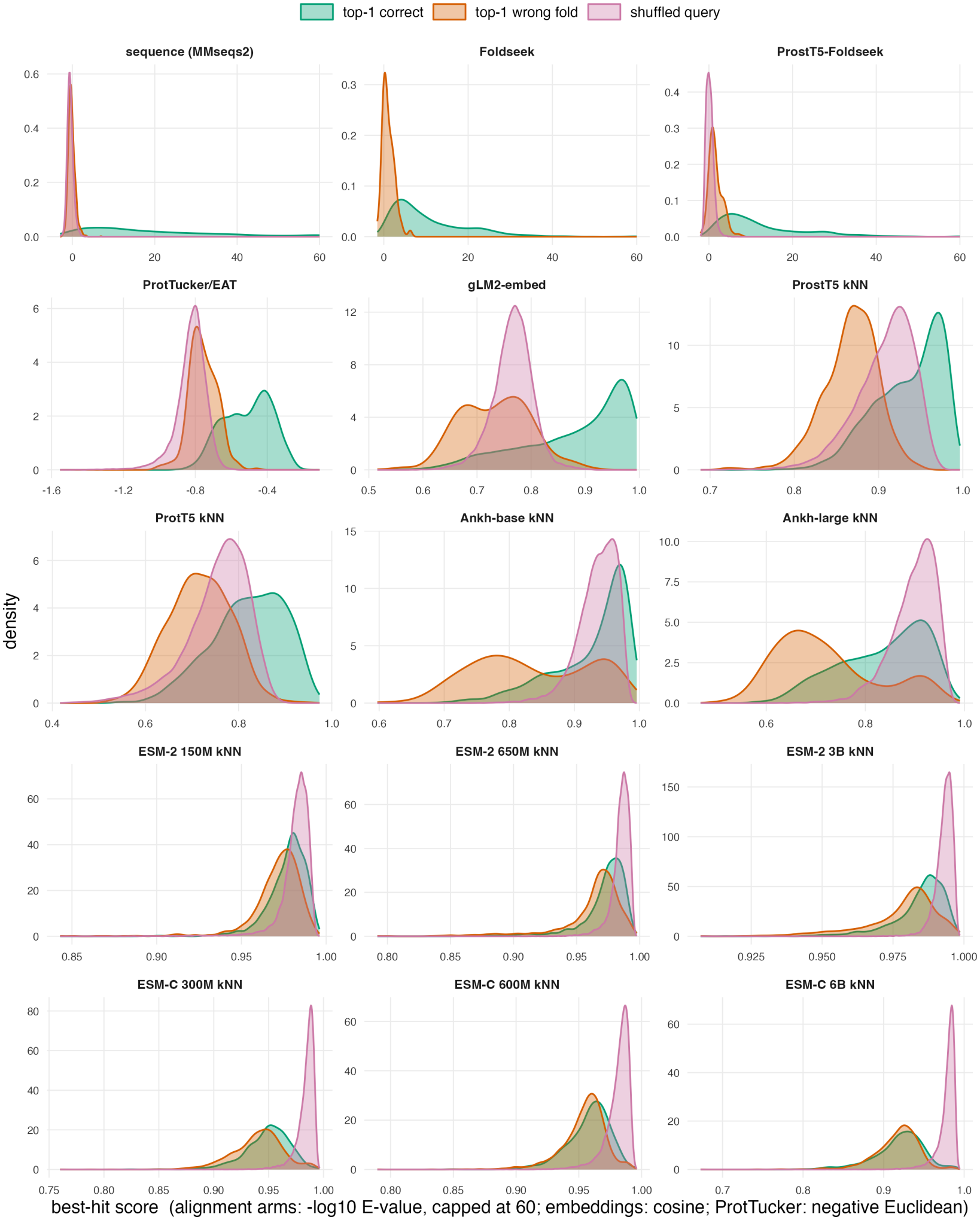
Density plots of cosine similarities/E-values of all methods on the Test 2 dataset for queries whose top hit is correct, queries whose top hit is in a different fold, and shuffled queries. Note ProtTucker uses Euclidean distance, stored negated so higher-is-better uniformly. Note also that Foldseek cannot have shuffled queries (as it requires query structures).

**Figure S4:**
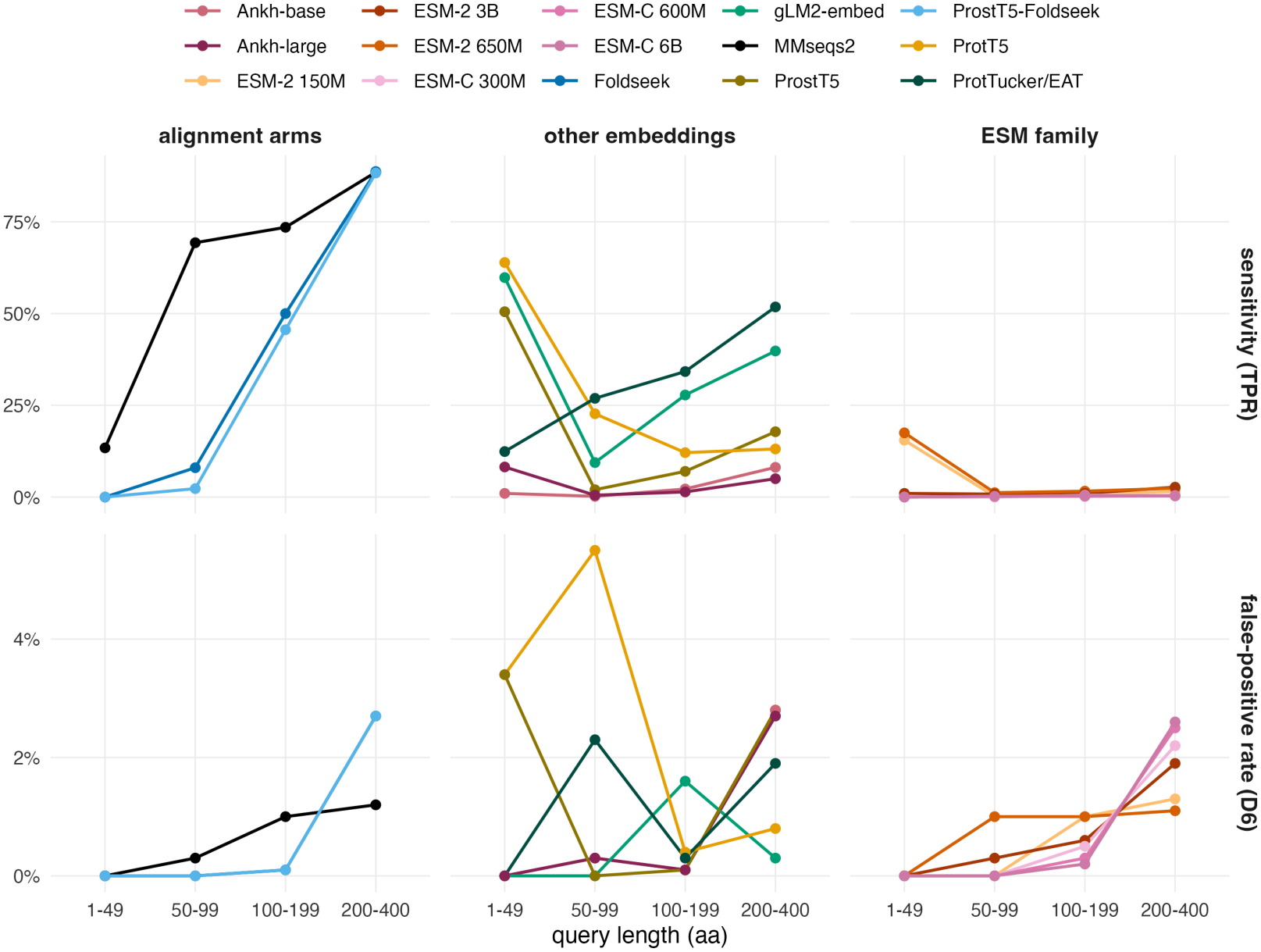
Sensitivity (TPR at 1% FPR calibrated across all queries) on Real Remote Homology and False Positive Rate of Natural Hard Negatives (D6) for Test 1 queries, stratified by length, across all tested methods. Note the 1–49 AA bin has only 97 positives and 29 negatives, and its boundary was chosen after inspecting the data, so it is exploratory.

**Figure S5:**
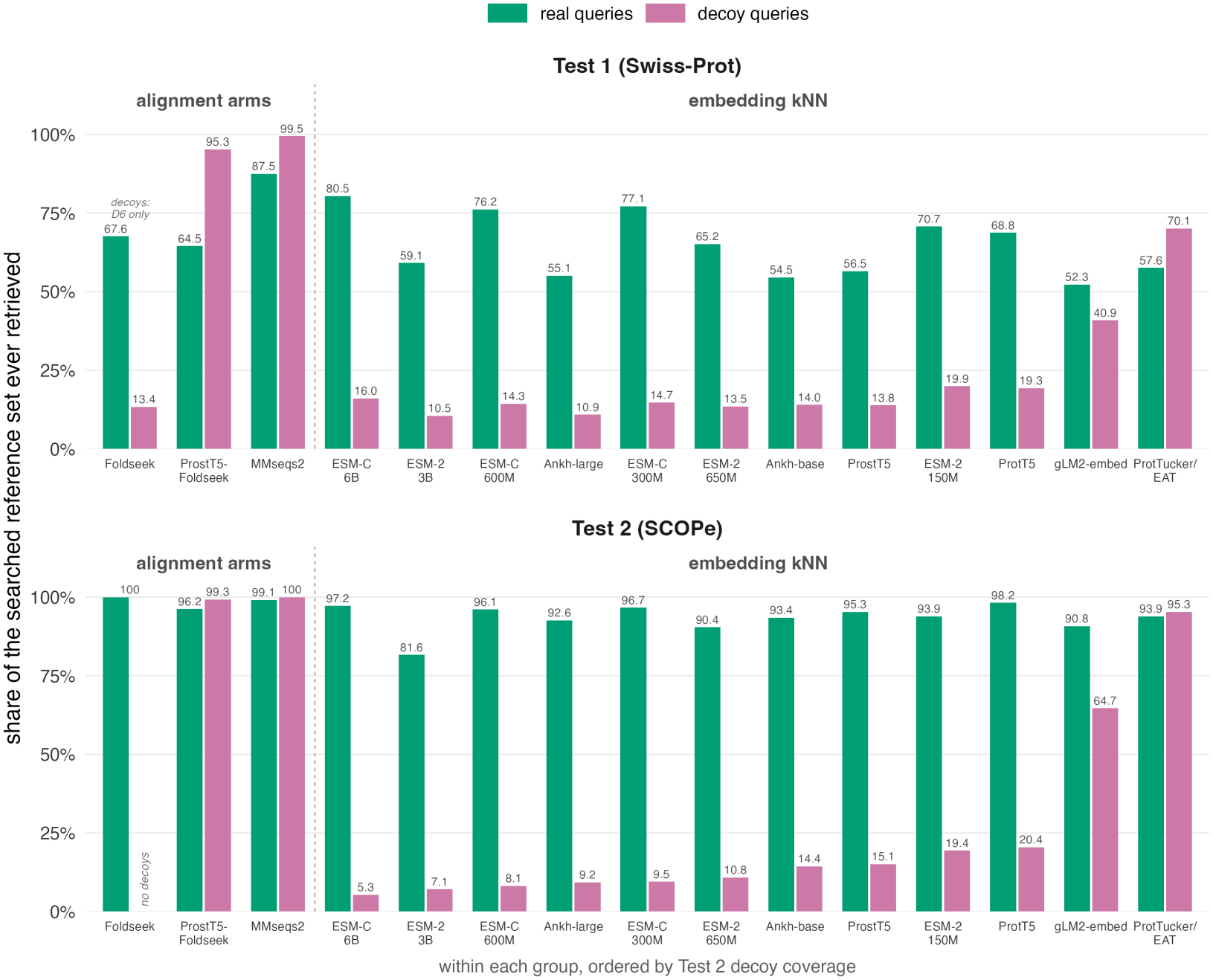
Coverage calculated as the fraction of reference entries appearing in any query’s top-50 list on Test 1 and Test 2 datasets across all methods for real remote homolog queries (green) and shuffled decoys (purple). For ProstT5-Foldseek, Foldseek and MMseqs2, this was calculated as the top 50 hits ranked by ascending E-value. Note Foldseek has no SCOPe decoys (as shuffling is not possible for query structures), and similarly only has D6 hard real negatives for Test 1.

**Figure S6:**
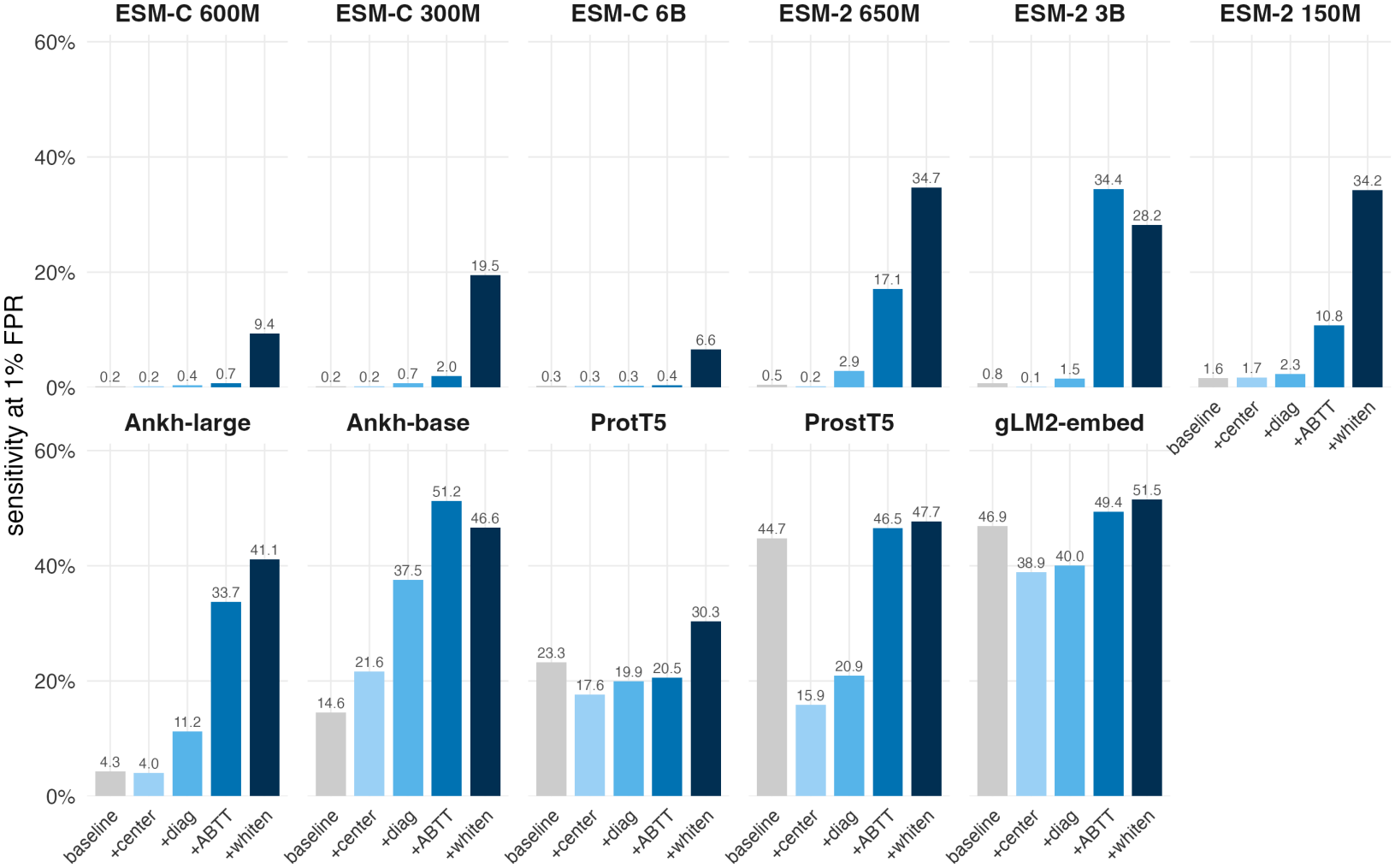
Sensitivity at 1% FPR for Test 2 (SCOPe) across all embedding models for the four correction methods fit on the reference database. Baseline cosine similarity (same as in Figure 1) is presented in grey, with ‘+center’ meaning Centering, ‘+diag’ meaning diagonal whitening, ‘+ABTT’ meaning All-but-the-top and ‘+whiten’ meaning ZCA whitening. Note ProtTucker is omitted as it is a supervised head rather than a raw pLM.

**Figure S7:**
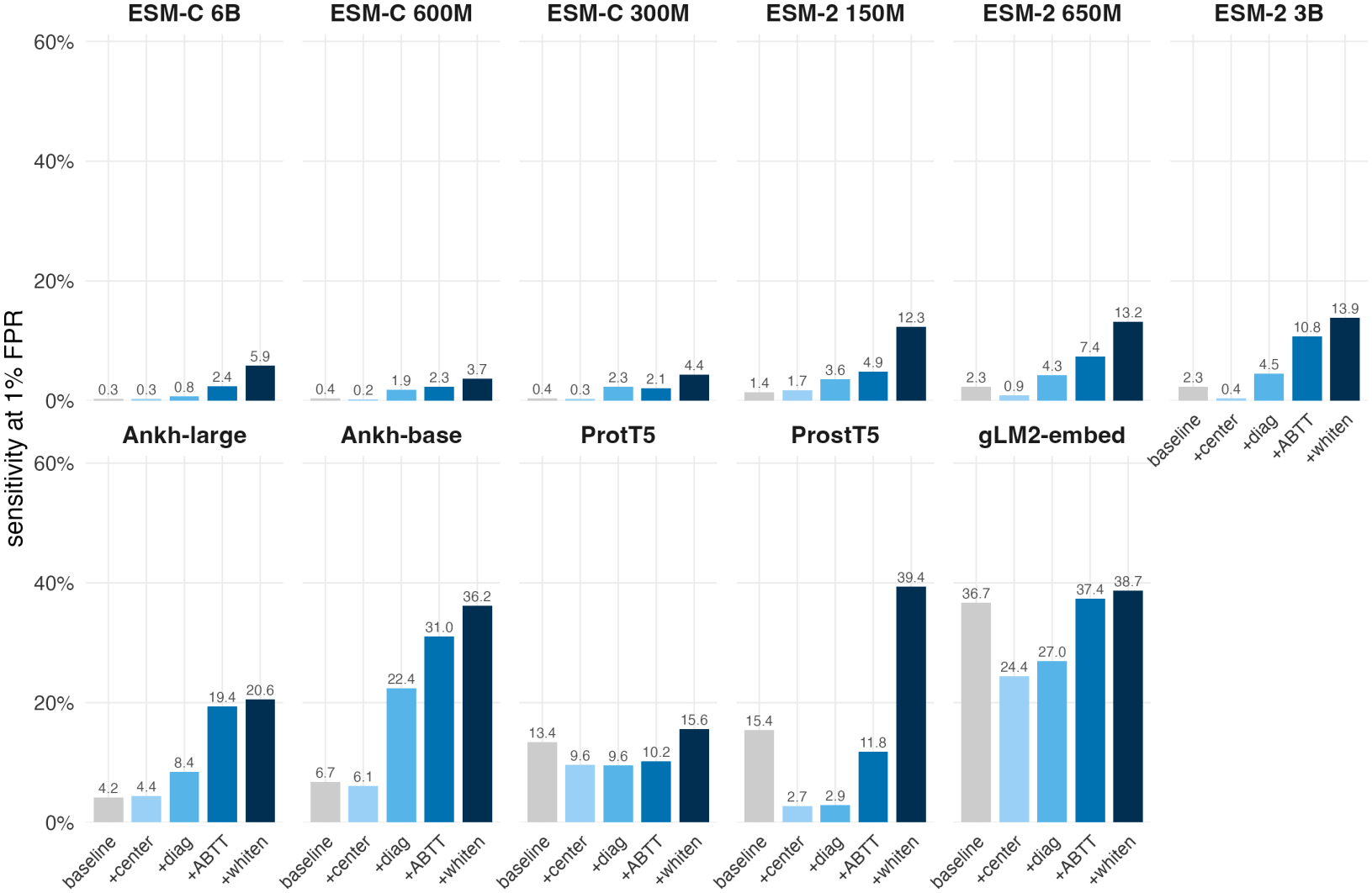
Sensitivity at 1% FPR for Test 1 (Swiss-Prot) across all embedding models for the four correction methods fit on the reference database. Baseline cosine similarity, with the threshold fitted once on the full data (Figure 1 shows bootstrap means), is presented in grey, with ‘+center’ meaning Centering, ‘+diag’ meaning diagonal whitening, ‘+ABTT’ meaning All-but-the-top and ‘+whiten’ meaning ZCA whitening. Note ProtTucker is omitted as it is a supervised head rather than raw pLM.

**Figure S8:**
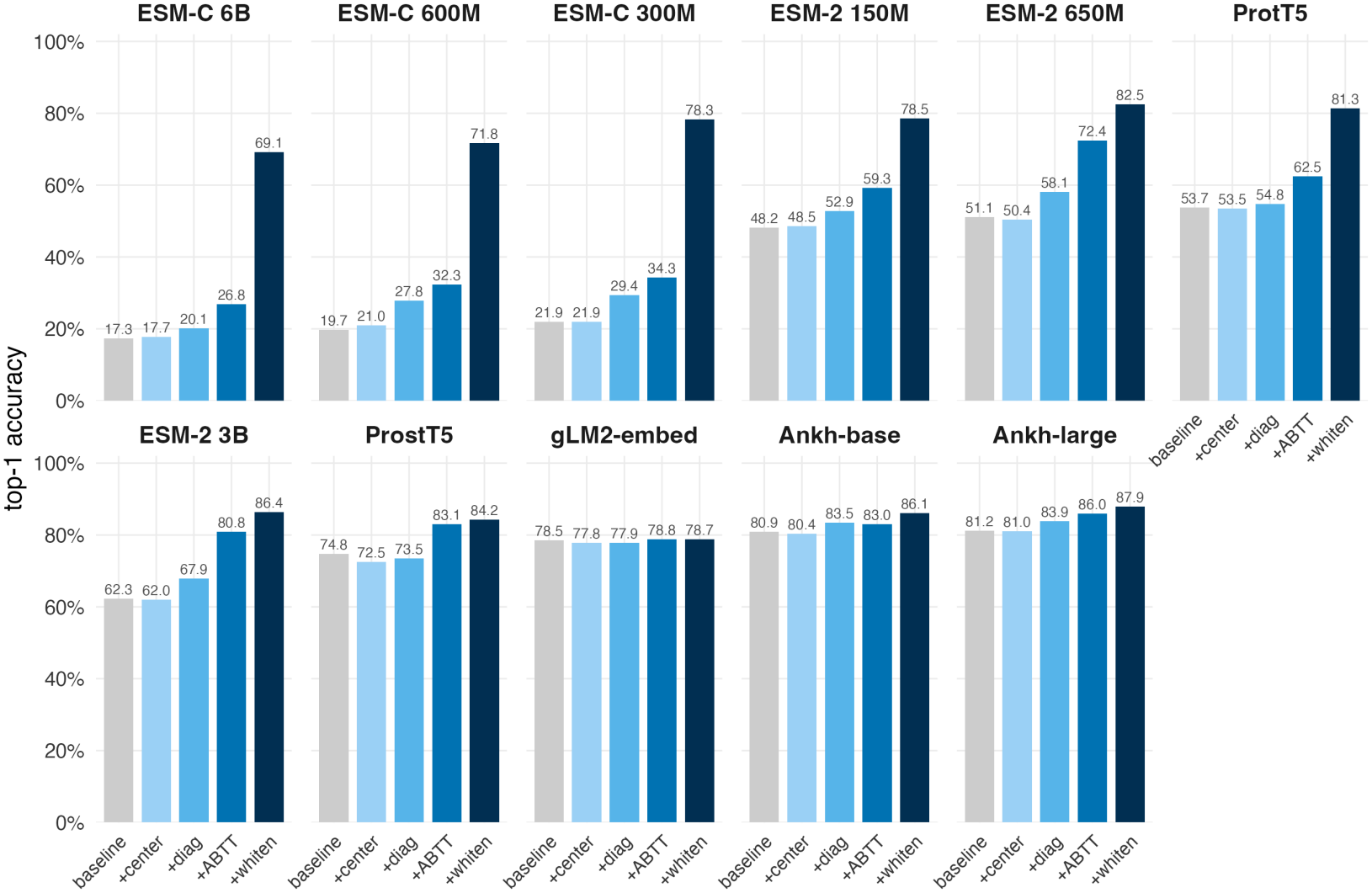
Threshold-free Top-1 retrieval for the four correction methods fit on the Test 1 reference database. Baseline cosine similarity is presented in grey, with ‘+center’ meaning Centering, ‘+diag’ meaning diagonal whitening, ‘+ABTT’ meaning All-but-the-top and ‘+whiten’ meaning ZCA whitening. Note ProtTucker is omitted as it is a supervised head rather than raw pLM.

**Figure S9:**
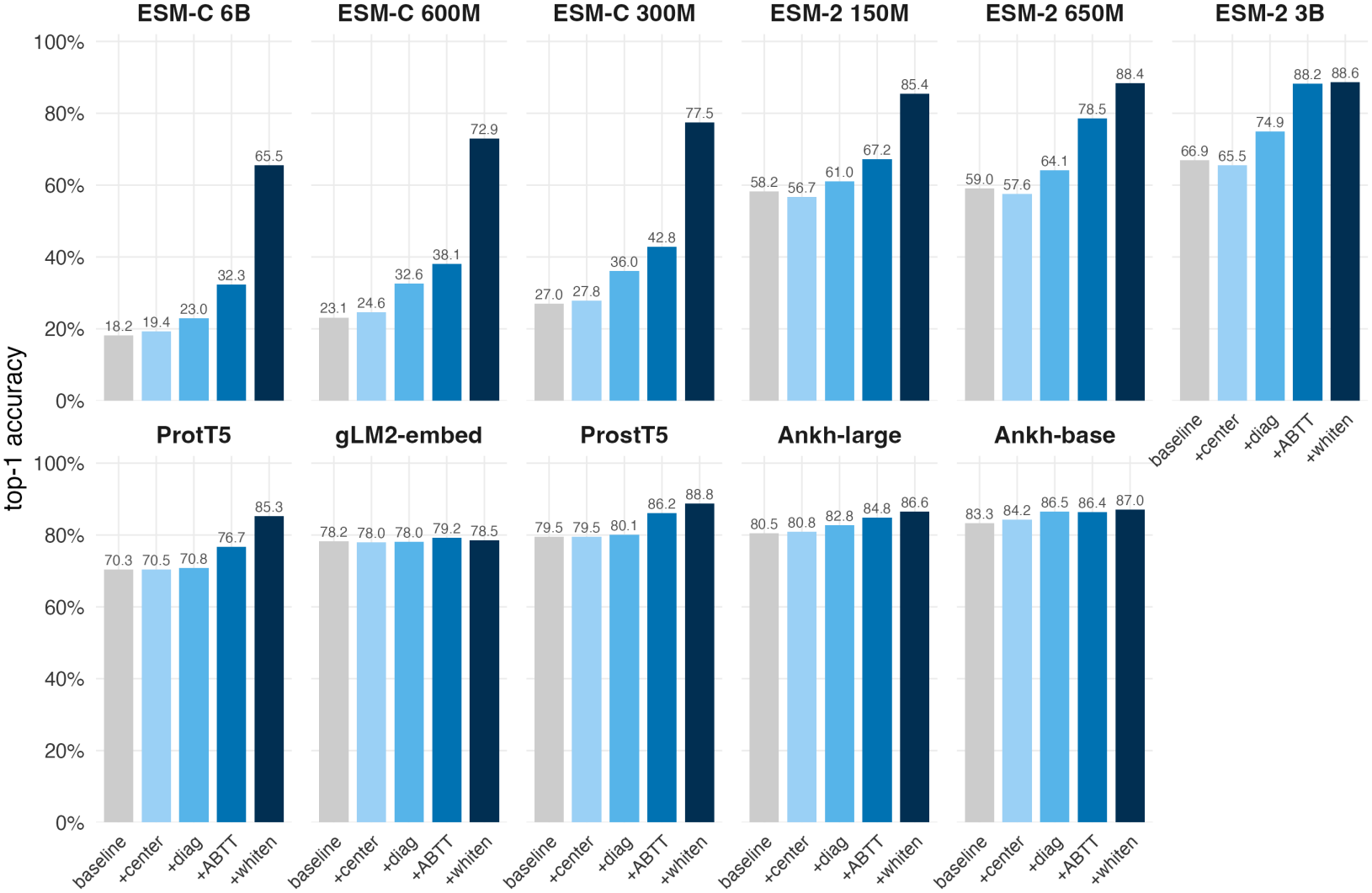
Threshold-free Top-1 retrieval for the four correction methods fit on the Test 2 reference database. Baseline cosine similarity (same as in Figure 3) is presented in grey, with ‘+center’ meaning Centering, ‘+diag’ meaning diagonal whitening, ‘+ABTT’ meaning All-but-the-top and ‘+whiten’ meaning ZCA whitening. Note ProtTucker is omitted as it is a supervised head rather than raw pLM.

**Figure S10:**
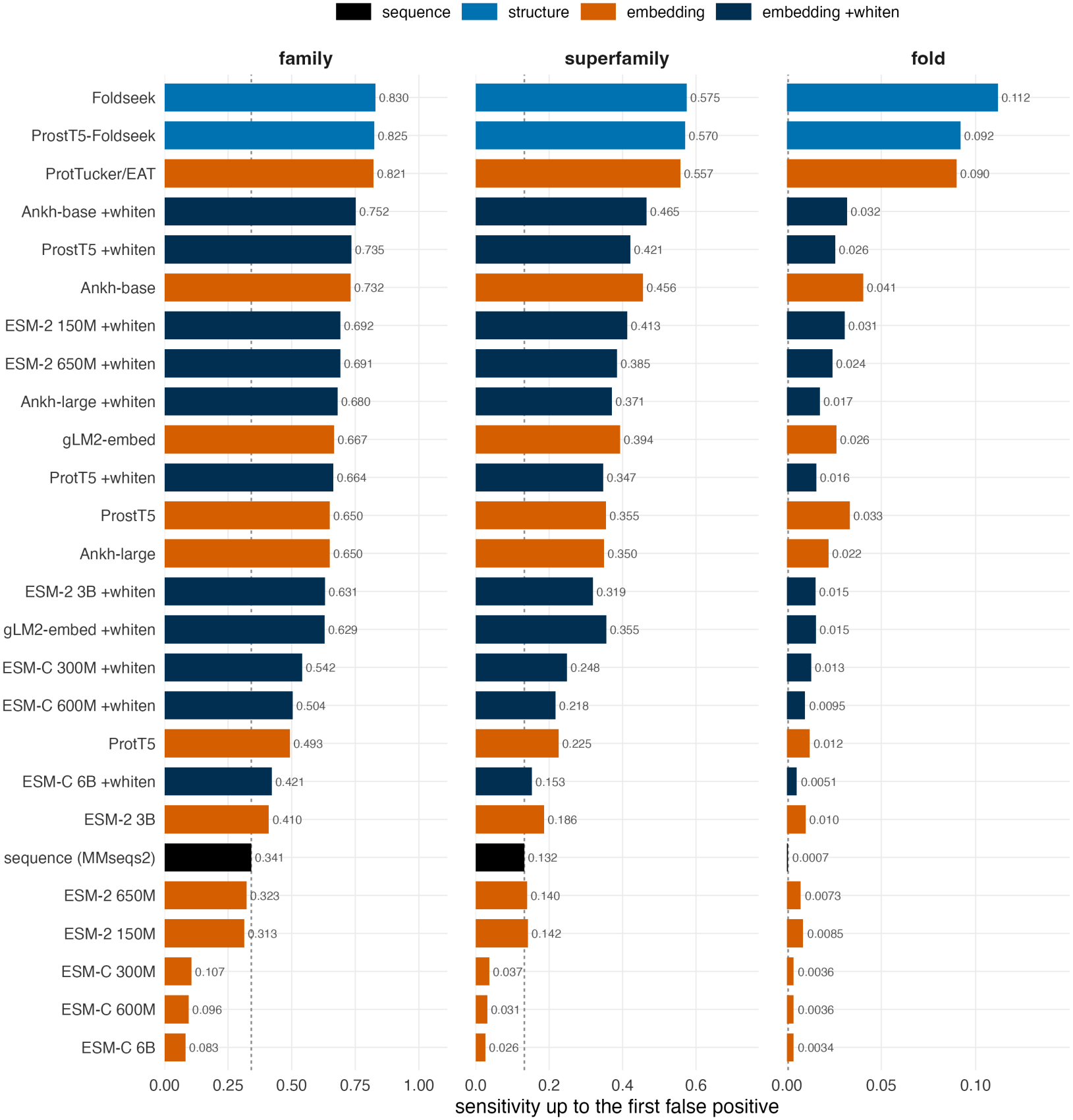
SCOPe 2.08 Foldseek benchmark (Test 3). The dashed line at each level (Family, Superfamily, Fold) indicates the MMseqs2 sequence baseline. Embedding-based models are indicated in orange, with ZCA-whitened versions in dark blue (excluding ProtTucker). As explained elsewhere, ProtTucker is expected to perform well, as it was explicitly finetuned on protein domain structures overlapping with the test set.

**Figure S11:**
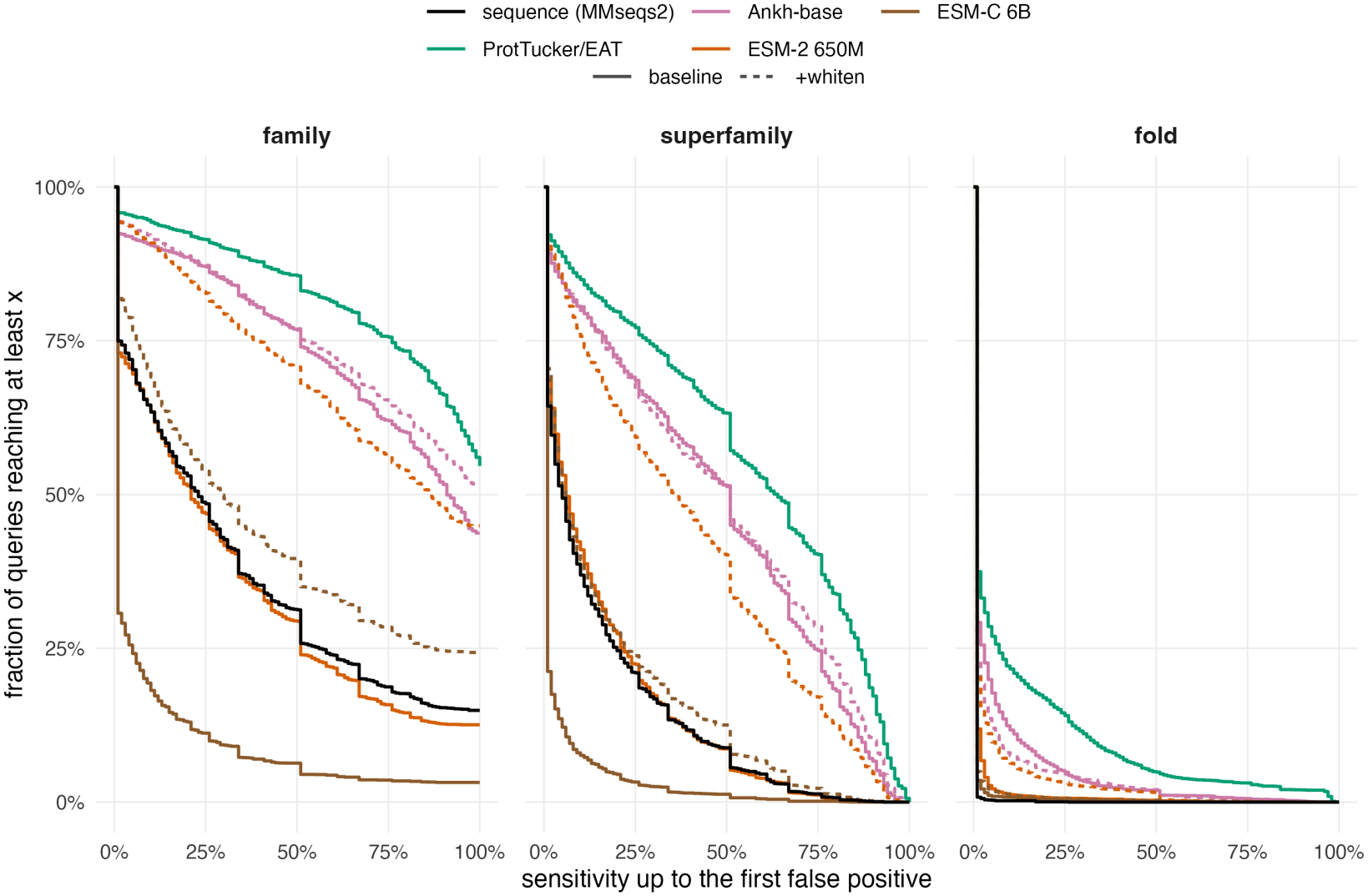
Cumulative curves for SCOPe 2.08 Foldseek benchmark (Test 3) at each level (Family, Superfamily, Fold). Only selected models were plotted in order to make the plot legible. Baseline pLM methods are drawn with solid lines, with ZCA-whitened methods indicated with a dashed line.

## Footnotes

1 More sophisticated sequence homology-based approaches, such as using position-specific scoring matrices (‘PSSMs’) and profile Hidden Markov models (‘pHMMs’) are also commonly used for annotation, but will not be considered in this manuscript as they require extra curation and are difficult to compare to embedding based methods.

2 Other practical issues that have hindered EAT’s widespread adoption include the enormous size of protein embedding databases compared with sequence and structure-based methods and the requirement for GPU-acceleration.

3 All pLMs tested would similarly include Swiss-Prot proteins as part of their pre-training corpus.

4 No layer sweep was run. This is an explicit limitation of our analysis. Intermediate layers may be better for some tasks [Joeres et al., 2026], though not necessarily [Vieira et al., 2025].

5 This reference database only approach matches the plausible real-world software deployment setting where the database would be shipped with the software.

6 Or indeed, as for some bacterial and bacteriophage annotation tasks, sometimes you do not even know that your query protein is a real protein at all [Cook et al., 2024, McNair et al., 2019].

7 Test-train leakage is nigh unavoidable as pLMs have been trained on essentially all well-understood proteins, and usually the functional or homolog labels have necessarily been created using some remote homology based inference method.

